# Cross-disease modeling of peripheral blood identifies biomarkers of type 2 diabetes predictive of Alzheimer’s disease

**DOI:** 10.1101/2024.12.11.627991

**Authors:** Brendan K. Ball, Jee Hyun Park, Elizabeth A. Proctor, Douglas K. Brubaker

**Affiliations:** Weldon School of Biomedical Engineering, Purdue University, West Lafayette, IN, USA; Department of Neurosurgery, Penn State College of Medicine, Hershey, PA, USA; Department of Pharmacology, Penn State College of Medicine, Hershey, PA, USA; Department of Biomedical Engineering, Penn State University, State College, PA, USA; Center for Neural Engineering, Penn State University, State College, PA, USA; Department of Engineering Science & Mechanics, Penn State University, State College, PA, USA; Center for Global Health & Diseases, Department of Pathology, School of Medicine, Case Western Reserve University School of Medicine, Cleveland, OH, USA; Blood Heart Lung Immunology Research Center, University Hospitals, Cleveland, OH, USA

**Keywords:** Type 2 diabetes, Alzheimer’s disease, transcriptomics, computational gene correlation analysis, cross-disease modeling, partial least squares discriminant analysis

## Abstract

Type 2 diabetes (T2D) is a significant risk factor for Alzheimer’s disease (AD). Despite multiple studies reporting this connection, the mechanism by which T2D exacerbates AD is poorly understood. It is challenging to design studies that address co-occurring and comorbid diseases, limiting the number of existing evidence bases. To address this challenge, we expanded the applications of a computational framework called Translatable Components Regression (TransComp-R), initially designed for cross-species translation modeling, to perform cross-disease modeling to identify biological programs of T2D that may exacerbate AD pathology. Using TransComp-R, we combined peripheral blood-derived T2D and AD human transcriptomic data to identify T2D principal components predictive of AD status. Our model revealed genes enriched for biological pathways associated with inflammation, metabolism, and signaling pathways from T2D principal components predictive of AD. The same T2D PC predictive of AD outcomes unveiled sex-based differences across the AD datasets. We performed a gene expression correlational analysis to identify therapeutic hypotheses tailored to the T2D-AD axis. We identified six T2D and two dementia medications that induced gene expression profiles associated with a non-T2D or non-AD state. Finally, we assessed our blood-based T2DxAD biomarker signature in post-mortem human AD and control brain gene expression data from the hippocampus, entorhinal cortex, superior frontal gyrus, and postcentral gyrus. Using partial least squares discriminant analysis, we identified a subset of genes from our cross-disease blood-based biomarker panel that significantly separated AD and control brain samples. Our methodological advance in cross-disease modeling identified biological programs in T2D that may predict the future onset of AD in this population. This, paired with our therapeutic gene expression correlational analysis, also revealed alogliptin, a T2D medication that may help prevent the onset of AD in T2D patients.

## INTRODUCTION

Type 2 diabetes (T2D) is a metabolic disease characterized by chronic hyperglycemia and insulin dysregulation that significantly elevates the risk for Alzheimer’s disease (AD) by more than 60%^1–3^. Alzheimer’s disease is an irreversible neurodegenerative disorder that gradually impairs memory and cognitive function. A recent large-scale longitudinal study found that individuals with an earlier onset of T2D were at higher risk of developing AD^4^. Other cohort studies^5,6^ reported similar results. In addition to the elevated risk of AD, T2D also contributes to other conditions such as hypertension^7^, neuroinflammation^8^, heart disease^9^, stroke^10^, and kidney disease^11^. As a result, the influence of T2D on other comorbidities further complicates our understanding of its impact on human health and the development of potential therapeutics for such conditions.

To understand this T2D-AD axis, previous studies examined how the onset of T2D influences the progression of AD^12^. Multiple studies reported insulin signaling impairment in T2D and AD^13,14^. The metabolic connection to AD^15^ also carries the T2D risk factor and is further amplified by the age^16^. Systemic low-grade inflammation in T2D progressively leads to downstream neuroinflammation and neuronal cell death, increasing the risk of AD^17–19^. Another study revealed altered gene expression levels in neurons, astrocytes, and endothelial cells in post-mortem brain tissue of T2D subjects, showing alterations to brain cells under diabetic conditions^20^.

Previous work from other groups implicates the blood-brain barrier (BBB) as a potential route that connects T2D^21^ and AD^22^. The BBB is a selective semipermeable membrane consisting of endothelial cells, pericytes, and astrocytes, which protects the brain from harmful substances and regulates the passage of immune cells and nutrients into the brain^23,24^. One large clinical study observed heightened BBB permeability in people with T2D and AD^25^. This progressive breakdown of the BBB in T2D and AD is associated with irregular vascular endothelial growth factor production, resulting in increased permeability across the BBB^25,26^. Other reports suggested that damage to endothelial cells in the cerebral blood vessels, indicated by elevated adhesion molecules, may contribute to this breakdown^25,27,28^. Therefore, chronic circulation of molecules produced under T2D conditions in the bloodstream may contribute to BBB breakdown and eventually enter the brain, contributing to the development of dementia and cognitive dysfunction.

A barrier to understanding how one disease influences another is that studies that simultaneously investigate multiple health conditions in humans are rare and difficult^29^. This challenge is compounded in chronic disorders like T2D and AD, where pathogenesis can precede diagnosis by decades^30^. To overcome this barrier, other groups have used differential expression analysis of transcriptomic data between T2D and AD but have fallen short in considering human heterogeneity, such as sex and age^31,32^. Another group integrated T2D and AD data using non-negative matrix factorization to identify shared genes across the blood of T2D and AD. While they identified dysregulated transcription factors shared across both diseases, they also did not account for confounding variables such as sex and age^33^. To overcome this challenge, we adapted Translatable Components Regression (TransComp-R), a computational approach initially developed to translate observations from pre-clinical animal disease models to human contexts^34–37^, to perform cross-disease modeling of human datasets to identify T2D biology predictive of AD.

In this work, we hypothesized that gene transcripts in T2D blood may predict and inform AD pathology. We tested this hypothesis via computational modeling of publicly available peripheral blood transcriptomics data of T2D and AD patients to determine if biomarkers in T2D blood could distinguish blood signatures in AD versus cognitively normal control groups. To identify potential therapeutics tailored to the T2D-AD axis, we employed a correlational analysis to identify candidate drugs that may impact AD development. Lastly, we assessed whether the blood-based biomarkers from our T2D-AD computational models could differentiate between AD and control samples in brain tissue transcriptomics data.

## RESULTS

### TransComp-R modeling separates AD and control subjects in T2D PC space

We acquired bulk-RNA seq T2D and microarray AD peripheral whole blood data from Gene Expression Omnibus (GEO). For the T2D dataset (GSE184050)^38^, we used the longitudinal baseline sample collection and information, including demographic variables of sex and age. Two separate cohorts of AD data were used in the model to test the predictability of T2D for AD. In both AD cohort 1 (GSE63060)^39^ and AD cohort 2 (GSE63061)^39^, we used AD and healthy control subjects. Using two separate cohorts ensured that the selected T2D PC’s would be robust **(Table 1)**.

**Table 1.**
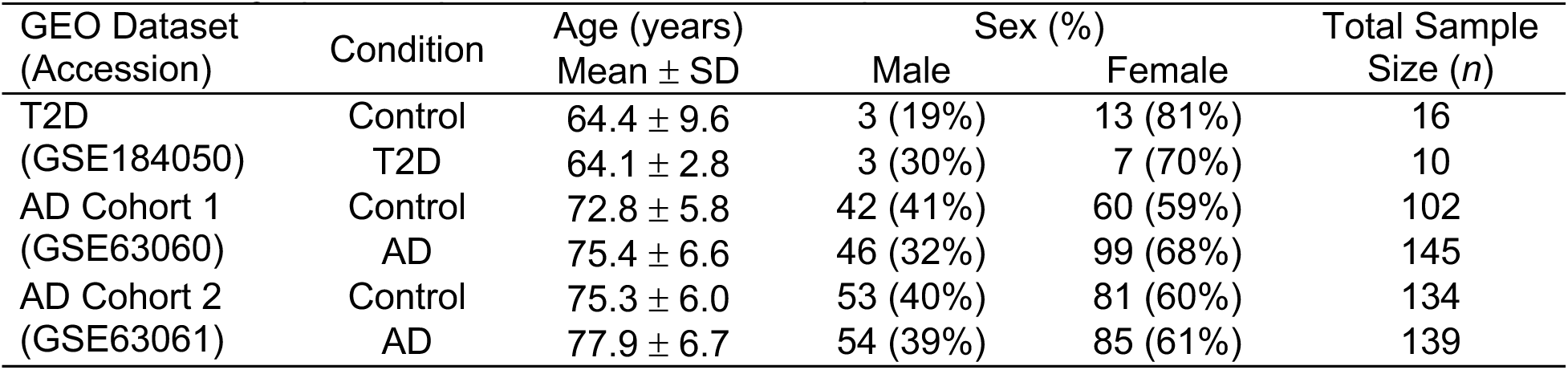
Demographics of processed human transcriptomic blood data across each data set.

| GEO Dataset<br>(Accession) | Condition | Age (years) | Sex (%) |  | Total Sample<br>Size (n) |
| --- | --- | --- | --- | --- | --- |
| | | Mean ± SD | Male | Female | |
| T2D<br>(GSE184050) | Control | 64.4 ± 9.6 | 3 (19%) | 13 (81%) | 16 |
| | T2D | 64.1 ± 2.8 | 3 (30%) | 7 (70%) | 10 |
| AD Cohort 1<br>(GSE63060) | Control | 72.8 ± 5.8 | 42 (41%) | 60 (59%) | 102 |
| | AD | 75.4 ± 6.6 | 46 (32%) | 99 (68%) | 145 |
| AD Cohort 2<br>(GSE63061) | Control | 75.3 ± 6.0 | 53 (40%) | 81 (60%) | 134 |
| | AD | 77.9 ± 6.7 | 54 (39%) | 85 (61%) | 139 |

We repurposed the TransComp-R to identify biological pathways dysregulated in T2D predictive of AD status. Cross-disease TransComp-R begins by matching shared genes across all datasets (**Fig. 1a**). We then projected the AD human samples into a principal component analysis (PCA) space constructed from the T2D data. We evaluate predictive power of T2D PCs for outcomes in AD by Least Absolute Shrinkage and Selection Operator (LASSO) feature selection and generalized linear model (GLM) regression (**Fig. 1b**). Using GSEA, we annotated the biological and therapeutic interpretations of the significant T2D PCs predictive of AD biology (**Fig. 1c**). We correlated differentially expressed genes from the drug list containing consensus signatures from the Library of Integrated Network-based Cellular Signatures (LINCS) database to the loadings of the T2D PCs predictive of AD. This method links drug regulation of genes associated with healthy states vs AD or T2D with drug response signatures to identify therapeutic hypotheses.

**Figure 1.**
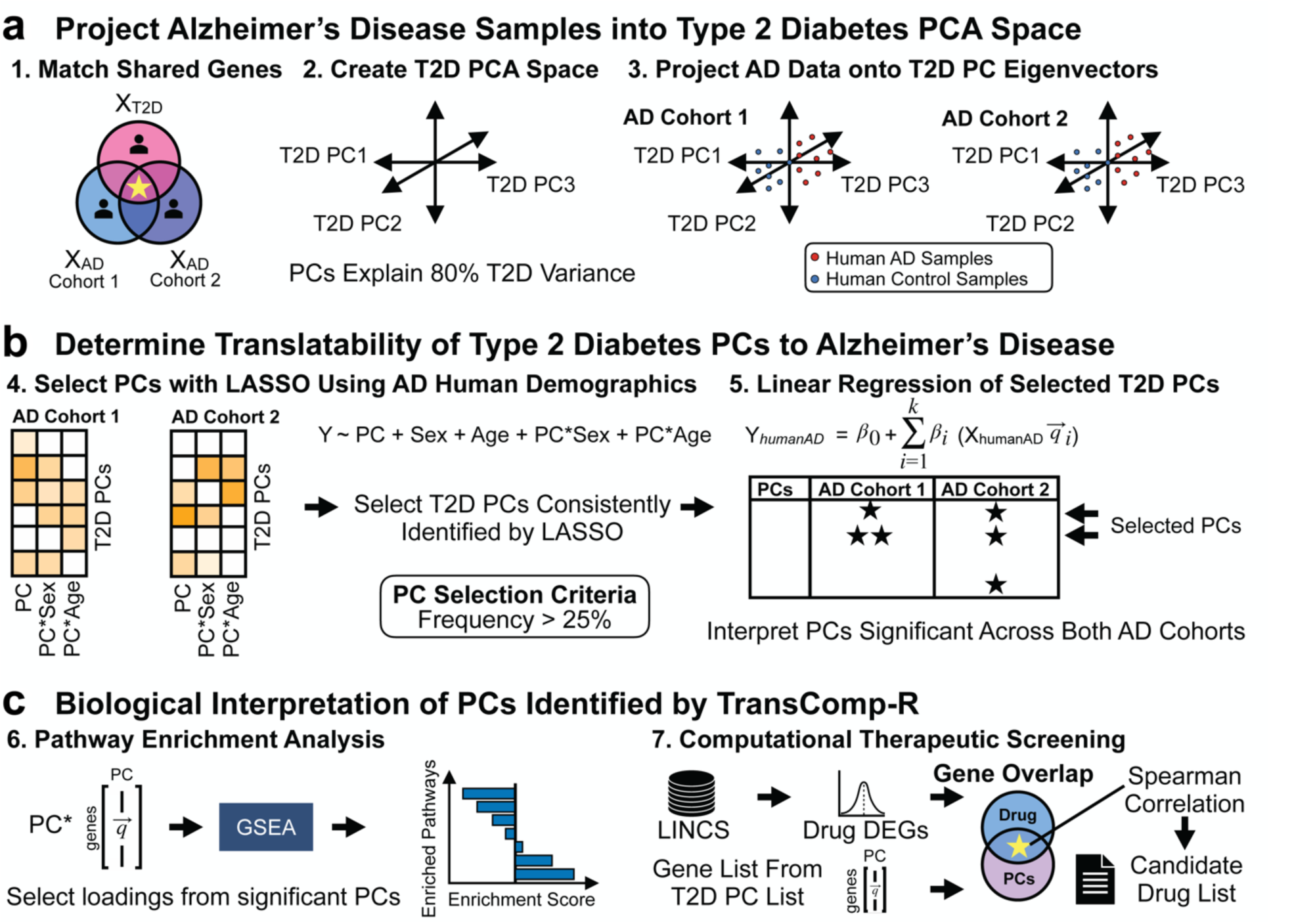
Workflow of TransComp-R. **(a)** Genes across T2D and AD are selected for analysis. Each AD cohort is individually projected into the T2D PCA space to combine the two diseases. **(b)** PC translatability from T2D to AD is determined by running a GLM regression against AD outcomes using PCs consistently selected across each AD cohort. **(c)** Pathway enrichment analysis is performed on the loadings of significant PCs to identify enriched biological pathways. Potential therapeutic candidates are then identified using a correlation analysis framework.

We matched 11,455 genes across the T2D and AD datasets and constructed the PCA space of the T2D and control samples. To prevent overfitting, we selected thirteen PCs for a cumulative explained variance of 80% for the TransComp-R model (**Supplementary Fig. S1**). Each AD cohort was separately projected onto the T2D PCs, such that we constructed two cross-disease models: T2D with AD cohort 1 and T2D with AD cohort 2.

We quantified how the variance captured by the T2D PCs explained the variation in human AD. To determine the cross-disease relevance of the T2D PCs to the variance of the AD data, we visualized each of the thirteen T2D PCs, comparing the variance explained in the T2D and AD data (**Fig. 2a**). When comparing the translatability of T2D PCs in AD cohort 1 and 2, we found T2D PC1, PC2, and PC3 had higher explained variance in Alzheimer’s disease data relative to the other T2D PCs 4-13, showing that T2D PCs1-3 have highest potential for translation of biology between T2D and AD.

**Figure 2.**
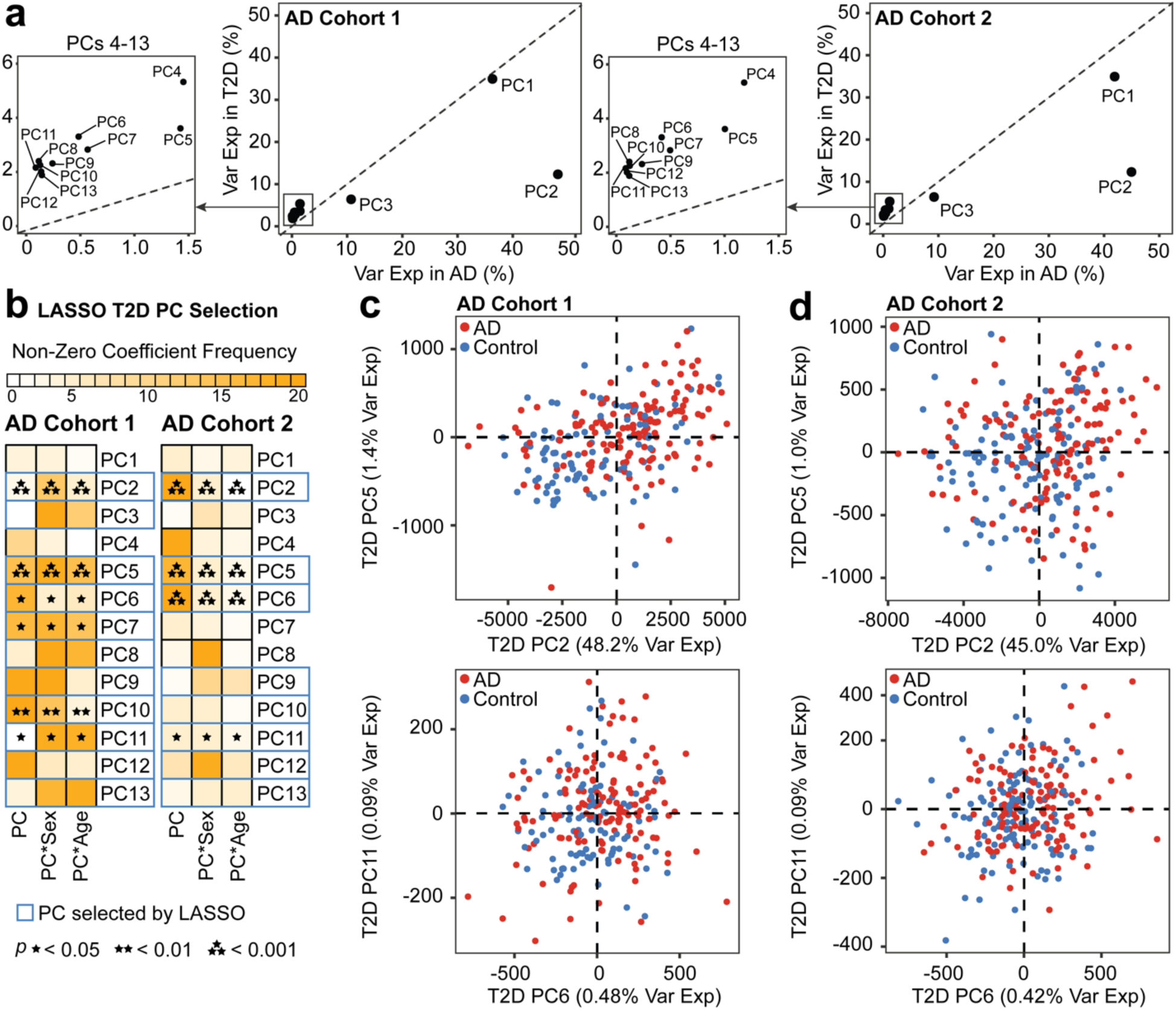
TransComp-R identifies T2D PCs predictive of AD outcomes. **(a)** AD PCs were separated by cohort, with variance explained in AD **(b)** Selection of PCs using a LASSO model incorporating sex and age demographics from the AD datasets. The model was run across twenty random rounds of ten-fold cross-validation. PCs consistently determined significant across both AD cohorts from the GLM regression were further analyzed. **(c)** Principal component plots of AD scores on selected T2D PCs separating AD and control outcomes in AD cohort 1 and **(d)** AD cohort 2. Each T2D PC is represented by the percent variance explained in AD.

We used LASSO to select the most relevant T2D PCs for predicting AD by regressing AD projections on T2D PCs, sex, and age from the AD cohort, with interaction effects of T2D PC with sex and age. From the LASSO model, several PCs (PC2, PC5-6, PC9-13) were selected across both AD cohorts (**Fig. 2b**). Despite the multiple number of PCs being consistently selected from LASSO, only T2D PC2, PC5, PC6, and PC11 fulfilled the selection criteria and discerned between AD and control groups in the GLM. The T2D PCs predictive of AD conditions were visualized for both AD cohort 1 (**Fig. 2c**) and AD cohort 2 (**Fig. 2d**). While the transcriptomic variation encoded on T2D PC2 and PC5 were able to distinguish between human AD and control groups, there was less distinguishable separation made by T2D PC6 and PC11. Among T2D PC2 and PC5, we selected T2D PC2 for deeper downstream interrogation due to the higher potential for T2D-to-AD translatability as quantified by the percentage of variance explained in AD (**Fig. 2a**).

### T2D and AD share pathways associated with metabolism, signaling pathways, and cellular processes

We employed GSEA to interpret the T2D PC2 gene loadings, which encoded transcriptomic variation between healthy and T2D subjects that predicted AD outcomes using both KEGG (**Fig. 3a**) and Hallmark (**Fig. 3b**) databases to gain a holistic insight into the genes loaded on T2D PC2.

**Figure 3.**
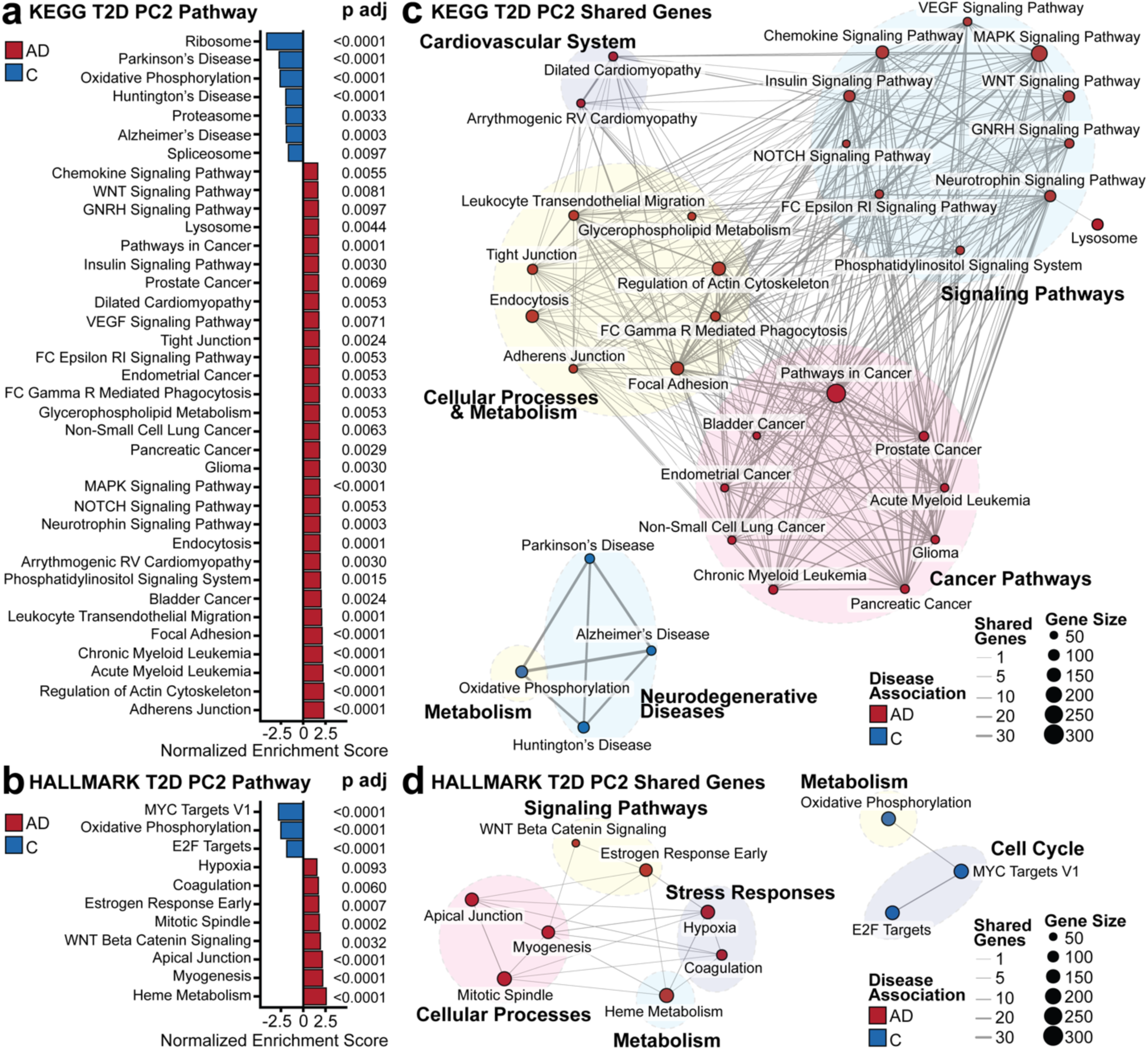
Pathway Enrichment Analysis. The transcriptomic variance separating AD and control subjects on T2D PC2 was interpreted with GSEA using the **(a)** KEGG and **(b)** Hallmark databases. Significantly enriched pathways were determined with a Benjamini-Hochberg adjusted *p* value less than 0.01. **(c)** Shared leading edge genes between biological pathways in the KEGG and **(d)** Hallmark pathways. The node size represents the number of genes contributing to the pathway from GSEA, whereas the edge size is the number of shared genes between each biological pathway. Missing pathways signified that there were no shared genes with other pathways.

We organized the enriched pathways into themes to determine if neighboring pathways were due to the overrepresentation of shared genes for both the KEGG (**Fig. 3c**) and Hallmark (**Fig. 3d**) databases. In the AD-associated pathways from KEGG, we identified enriched pathway themes, such as the cardiovascular system, signaling pathways, cellular processes and metabolism, and cancer pathways. In the control group, we found pathways associated with neurodegenerative diseases and metabolism. From Hallmark, pathways enriched in AD associations included signaling pathways, cellular processes, metabolism, and stress response, with metabolism and cell cycle pathways enriched in controls.

### T2D PC2 identifies gene expression changes with predictive ability across sex and disease conditions in two AD cohorts

We compared the average log_2_ fold change of the 11,455 shared genes for disease and control groups to identify trends in the regulation of genes across diseases. In both AD cohorts and T2D, there were decreases in gene expression including *COX7C*, *NDUSF5*, *NDUFA1*, *RPL17*, *RPL23*, *RPL26*, *RPL31*, and *TOMM7* (**Fig 4a**), genes responsible for mitochondrial and ribosomal functions. *COX7C*, *NDUSF5*, and *NDUFA1* are active in the electron transport chain function in the inner mitochondrial membrane and *TOMM7* encodes for a subunit of the translocase of the outer mitochondrial membrane. Ribosomal protein L genes such as *RPL17*, *RPL23*, *RPL26*, and *RPL31* play a role in forming structures of ribosomes and regulating ribosome function.

**Figure 4.**
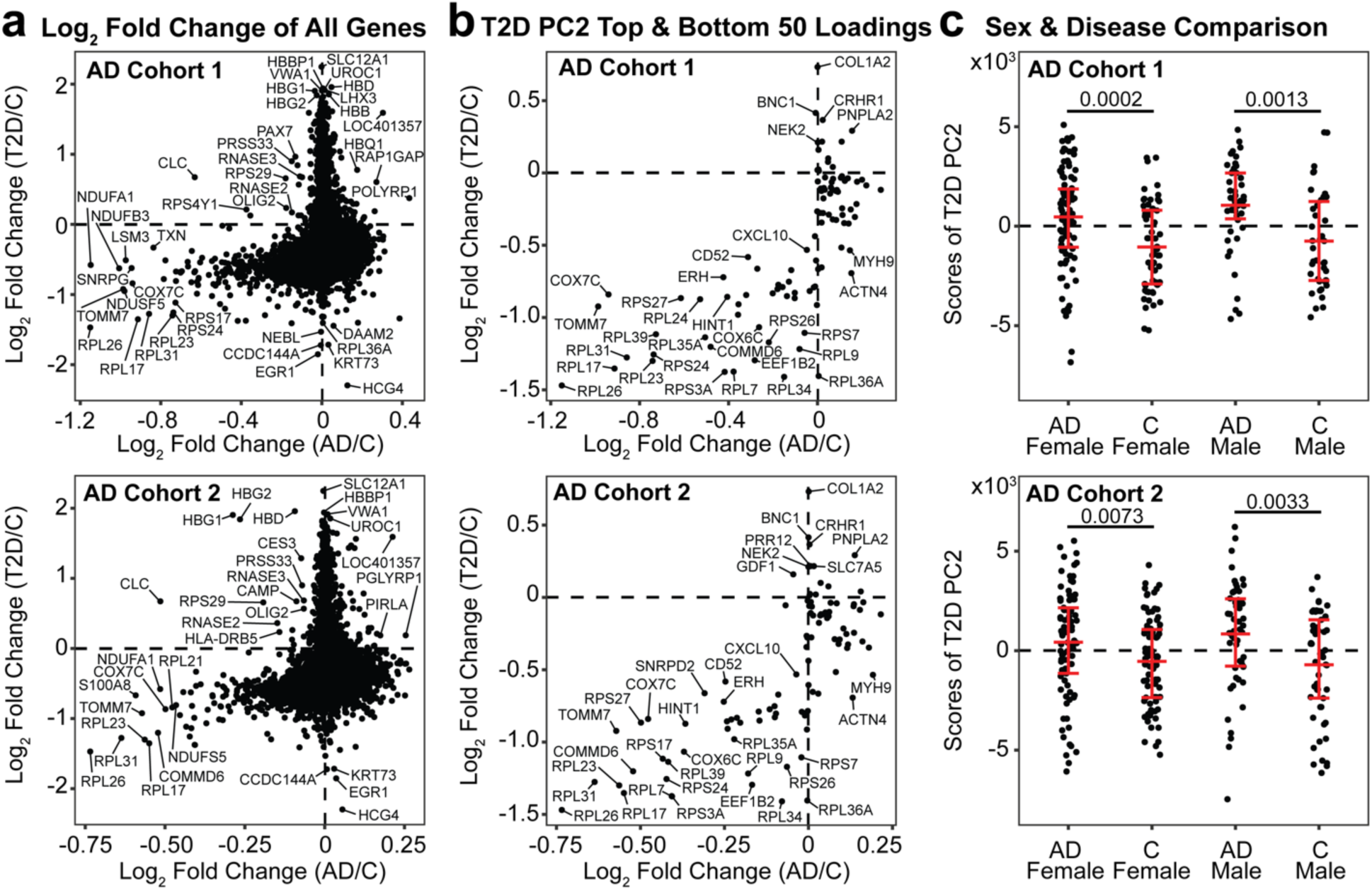
Comparison of global gene expression and AD-predictive T2D PCs. **(a)** AD and T2D log_2_ fold change plot of all shared 11,455 genes **(b)** AD and T2D log_2_ fold change plot filtered by gene expressions with the top 50 and bottom 50 loadings of T2D PC2. **(c)** Scores of T2D PC2 separated by sex and disease condition. A Mann-Whitney test adjusted by Benjamini-Hochberg was used to determine statistical significance. The distribution of the data is annotated by the mean and interquartile ranges.

We next tested to see if the top 50 and bottom 50 gene loadings from T2D PC2 could capture the cross-disease trends of the total transcriptome. We visualized the filtered gene with AD and T2D fold changes and observed a similar trend such that multiple genes were downregulated in both AD and T2D conditions (**Fig. 4b**). Among those consistently downregulated in AD and T2D, genes related to ribosomal proteins (RPL and RPS) were present. These 100 genes also distinguished between control and AD subjects (**Supplementary Fig. S2**).

Finally, we evaluated T2D PC2’s ability to stratify sex and disease characteristics in AD. We identified significant sex-based differences across AD and control in both cohorts. In AD cohort 1, we found that the female and male groups, each separated by AD and control, were significantly different by the variation captured by T2D PC2, with adjusted *p* values of 0.0002 and 0.0013, respectively (**Fig. 4c**). Similarly, in AD cohort 2, there was significance in disease separation for both females and males, with adjusted *p* values of 0.0073 and 0.0033, respectively (**Fig. 4c**). Comparing the scores of T2D PC2 by disease condition only, we found significance in both AD cohort 1 (*p* = 2.000×10^−7^) and AD cohort 2 (*p* = 9.078×10^−5^).

### Identification of drug perturbation signatures associated with PC2 T2D-AD signatures

We developed a correlation analysis to identify therapeutic candidates associated with the T2D PC2 predictive of AD. We used the Library of Integrated Network-Based Cellular Signatures (LINCS) Consensus Signatures, a dataset containing 33,609 drugs with their respective post-treatment gene expression profiles summarized as a “characteristic direction” (CD) coefficient^40^. Of the 33,609 drugs in the LINCS database, 3,161 remained after we filtered out duplicates and drugs without known targets. We compared the CD coefficient values of genes affected by each drug to the gene loadings on T2D PC2 using Spearman’s correlation. We hypothesized a drug could be therapeutic for T2D/AD risk based on the correlation directionality, where negative ρ values were interpreted as inducing profiles associated with a non-T2D or non-AD state and positive ρ values associated with a T2D or AD disease state.

We identified 1,262 drugs significantly correlated with the loadings in T2D PC2 (**Fig. 5a**). Drugs associated with a non-T2D and non-AD gene expression profile included dienestrol, BW-180C, T-0156, alogliptin, and roflumilast (**Supplementary Table S1**). Dienestrol had the most negative correlation coefficient of −0.5059 and is an estrogen receptor agonist used to treat vaginal pain by targeting *ESR1*. T-0156 (*PDE5A*) and roflumilast (*PDE4A*, *PDE4B*, *PDE4C* and *PDE4D*) are both phosphodiesterase inhibitors. We also identified a prototypical delta opioid receptor agonist (BW-180C) and a T2D prescription medication (alogliptin), which targets *OPRD1* and *DPP4*, respectively. Conversely, drugs associated with gene expression of a T2D or AD disease state included antagonists such as wortmannin (*PI3K* inhibitor), proglumide (*CCK* receptor antagonist), GR-127935 (serotonin receptor antagonist), homatropine-methylbromide (acetylcholine receptor antagonist), and phenacemide (sodium channel blocker).

**Figure 5.**
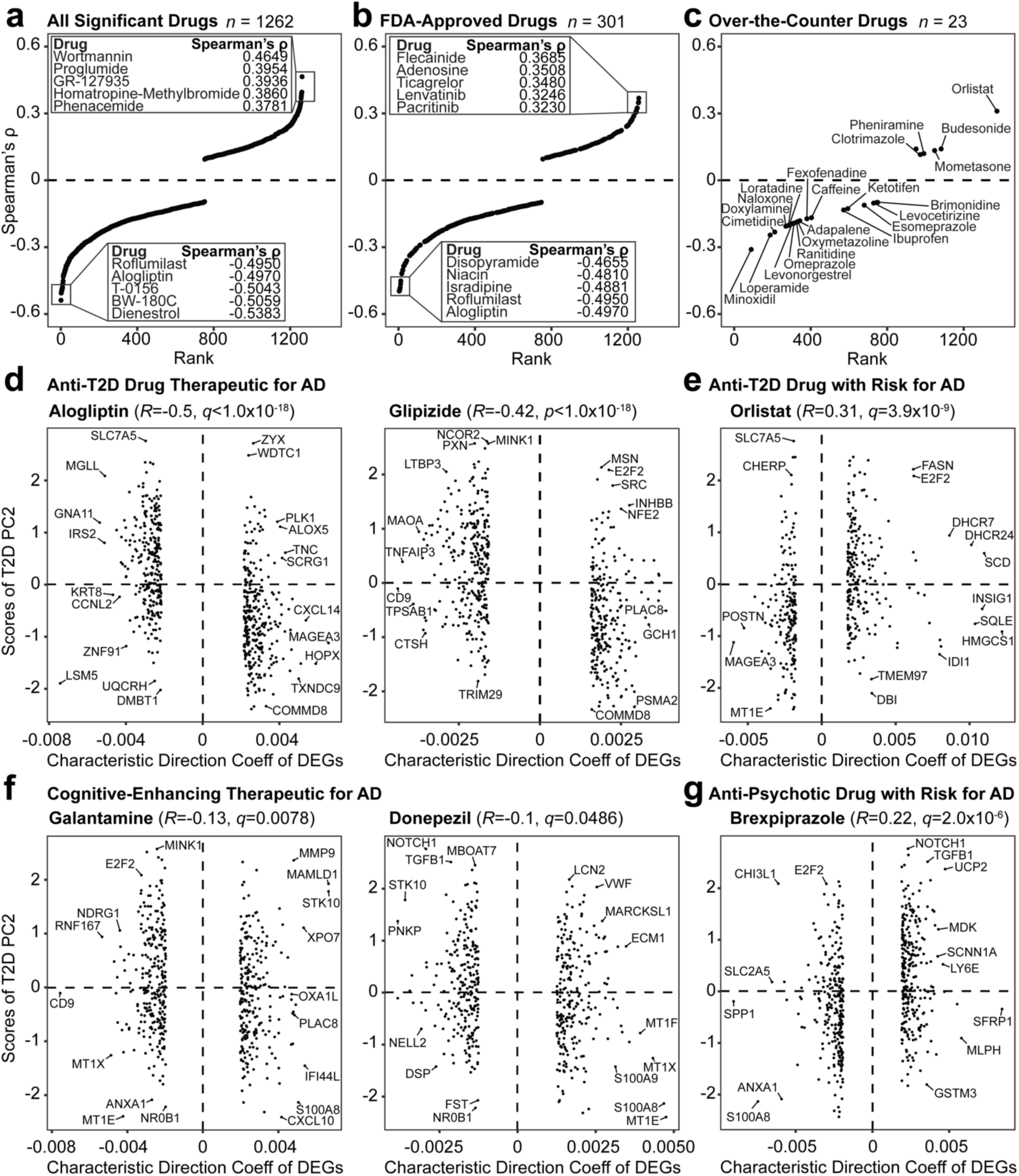
Computational gene expression correlational analysis. **(a)** All significant drugs identified from the LINCS database. Drugs filtered by **(b)** FDA approval status and **(c)** over-the-counter drugs. **(d)** FDA-approved T2D drugs (alogliptin and glipizide) associated with control group signatures. **(e)** FDA-approved T2D drug (orlistat) associated with genes upregulated in AD. **(f)** FDA-approved medications for cognitive-enhancement (galantamine and donepezil). **(g)** FDA-approved drug (brexpiprazole) with signatures correlated to genes elevated in AD.

To filter drugs tested for safety and efficacy, we referenced the Food and Drug Administration (FDA) Orange Book for FDA-approved and over-the-counter drugs (June 2024 version)^41^. We identified 301 FDA-approved drugs in our original significant 1,262 (**Fig. 5b**), and of these, 23 were approved for over-the-counter use (**Fig. 5c**). Among the FDA-approved drugs, alogliptin and roflumilast were among the most negative correlation coefficients. Other medications with negative coefficients associated with a non-T2D or AD state were isradipine, used for hypertension (*CACNA1S*, *CACNA1C*, *CACMA1F*, *CACMA1D*, and *CACMA2D1* targets), niacin used for vitamin B (*HCAR2* and *HCAR3* targets), and disopyramide used for irregular heartbeats (*SCN5A* gene target) (**Supplementary Table S2)**. Among medications with top positive coefficients associated with AD and T2D, we identified two anti-cancer drugs (pacritinib and lenvatinib), a blood thinner (ticagrelor), and two anti-arrhythmic drugs (adenosine and flecainide).

The most negative coefficients for over-the-counter drugs were vasodilators, opioid receptor targets, and histamine receptor drugs (**Supplementary Table S3**). Minoxidil had the most negative correlation coefficient (−0.3101) and is a hypertension medication that targets *KCNJ8*, *KCNJ11*, and *ABCC9*. Loperamide (opioid receptor agonist), used for diarrhea, targets *OPRM1* and *OPRD1*, while naloxone (opioid receptor antagonist), used for opioid overdose, affects *OPRK1*, *OPRM1*, and *OPRD1*. We also identified two histamine receptor antagonists, cimetidine and doxylamine, which targeted *HRH2* and *HRH1*, respectively. The most positively correlated medications that induced disease gene signatures included orlistat, a lipase inhibitor used for weight loss and T2D, had the greatest coefficient of 0.3104 (*LIPF*, *PNLIP*, *DAGLA*, and *FASN* targets). Other positive correlation, T2D-AD associated drugs included budesonide (corticosteroid for Crohn’s disease) and mometasone (steroid for skin discomfort), both of which are glucocorticoid receptor agonists with the target of *NR3C1*. Other medications among the most positively correlated included clotrimazole (cytochrome p450 inhibitor) and pheniramine (histamine receptor antagonist), which targeted *KCNN4* and *HRH1* respectively.

We compared the FDA-approved drugs to MedlinePlus and First Databank for any medication currently used to treat T2D or cognitive-associated symptoms (**Supplementary Table S4**). Of the 301 FDA-approved drugs identified, we found ten medications for T2D and three with cognitive function associations (**Supplementary Table S5**). Among the medications used for T2D, glipizide (sulfonylurea), repaglinide (insulin secretagogue), and nateglinide (insulin secretagogue) targeted *KCNJ11* and *ABCC8*. The diabetes dipeptidyl peptidase inhibitors that target *DPP4,* included alogliptin, sitagliptin, and linagliptin. We also identified sodium/glucose co-transporter inhibitor empagliflozin (*SLC5A2*), the *PPAR* receptor antagonist pioglitazone, glucosidase inhibitor acarbose (*AMY2A*, *MGAM*, and *GAA*), and lipase inhibitor orlistat (*LIPF*, *PNLIP*, *DAGLA*, and *FASN*). Among medications commonly prescribed to improve cognitive function, we identified donepezil and galantamine, acetylcholinesterase inhibitors that target *ACHE* and *ACHE*/*BCH*E and brexpiprazole (*HTR2A*, *DRD2*, *HTR1A)*, a dopamine receptor partial agonist used for AD-associated agitation. Of these thirteen medications, empagliflozin, linagliptin, brexpiprazole, acarbose, and orlistat contained gene expression responses correlated to an AD or T2D condition. Nine medications were associated with a non-AD or non-T2D condition, which included alogliptin, glipizide, repaglinide, sitagliptin, pioglitazone, galantamine, nateglinide, and donepezil.

We selected the top two medications that associated with a non-disease state (T2D and cognitive-enhancing medication) and those associated with a disease state to compare the relationship of the drug DEGs and T2D PC2 scores. We found that alogliptin and glipizide, anti-T2D drugs had the most significant correlation magnitude among the six drugs, with a coefficient of −0.5 (*p* < 2.2×10^−16^) and −0.42 (*p* < 2.2×10^−16^), respectively (**Fig. 5d**). Orlistat had gene signatures most positively correlated with disease states (rho = 0.31, *p* = 2.9×10^−10^) (**Fig. 5e**). The signatures affected by cognitive medications galantamine (rho = −;0.13 *p* = 0.0028) and donepezil (rho = −;0.1 *p* = 0.024) had weaker correlations than the anti-T2D medication (**Fig. 5f**). Finally, we identified brexpiprazole, an anti-psychotic drug with a low positive correlation coefficient of 0.22 (*p* = 2.6×10^−7^) associated with T2D and AD disease status (**Fig. 5g**). Other FDA-approved T2D medications, with weaker correlations to a non-T2D or non-AD state included repaglinide, sitagliptin, pioglitazone, and nateglinide (**Supplementary Fig. S3**).

### Translation of T2D PC2 gene loadings to from AD blood to AD brain transcriptomics

Having identified biomarkers in T2D blood predictive of AD status, we assessed if the identified signature stratified AD from control patients in brain tissues. We acquired a human microarray dataset (GSE48350)^42,43^ profiling AD and control samples in multiple brain regions: hippocampus, entorhinal cortex (EC), superior frontal gyrus (SFG), and postcentral gyrus (PoCG). Potential age bias was reduced by excluding subjects younger than 55. The post-processed demographics separated by their respective brain region were summarized (**Table 2**).

**Table 2.** Demographic summary across four different processed human brain regions

| GEO Dataset<br>(GSE48350) | Condition | Age (years)<br>Mean ± SD | Sex (%) | | Total Sample<br>Size (n) |
| --- | --- | --- | --- | --- | --- |
|  |  |  | Male | Female |  |
| Hippocampus | Control | 82.0 ± 10.0 | 13 (52%) | 12 (48%) | 25 |
| | AD | 83.1 ± 8.5 | 9 (47%) | 10 (53%) | 19 |
| Entorhinal Cortex | Control | 80.7 ± 10.3 | 9 (50%) | 9 (50%) | 18 |
| | AD | 86.5 ± 5.5 | 7 (47%) | 8 (53%) | 15 |
| Superior Frontal<br>Gyrus | Control | 80.8 ± 10.3 | 12 (46%) | 14 (54%) | 26 |
| | AD | 87.1 ± 6.2 | 7 (33%) | 14 (67%) | 21 |
| Postcentral Gyrus | Control | 81.5 ± 10.4 | 11 (46%) | 13 (54%) | 24 |
| | AD | 85.0 ± 8.2 | 10 (40%) | 15 (60%) | 25 |

We matched genes in the AD brain dataset to the top 50 and bottom 50 genes from T2D PC2 (**Fig. 6a**) and matched 88 genes. We determined AD status-associated genes in each brain region via differential expression analysis (Benjamini-Hochberg adjusted Mann-Whitney test, *p* adjusted < 0.20). We first investigated the hippocampus brain tissue to identify genes from T2D-blood PC2 that could stratify AD and control groups in the brain. We identified 25 significant genes (adjusted *p* value < 0.20) and hierarchical clustering showed these 25 genes separated AD and control conditions in the hippocampus gene expression data (**Fig. 6b**). We used these genes to construct PLS-DA models to identify genes driving separation across the brain tissue samples of AD and control groups (**Fig. 6c, Supplementary Fig. S4)**.

**Figure 6.**
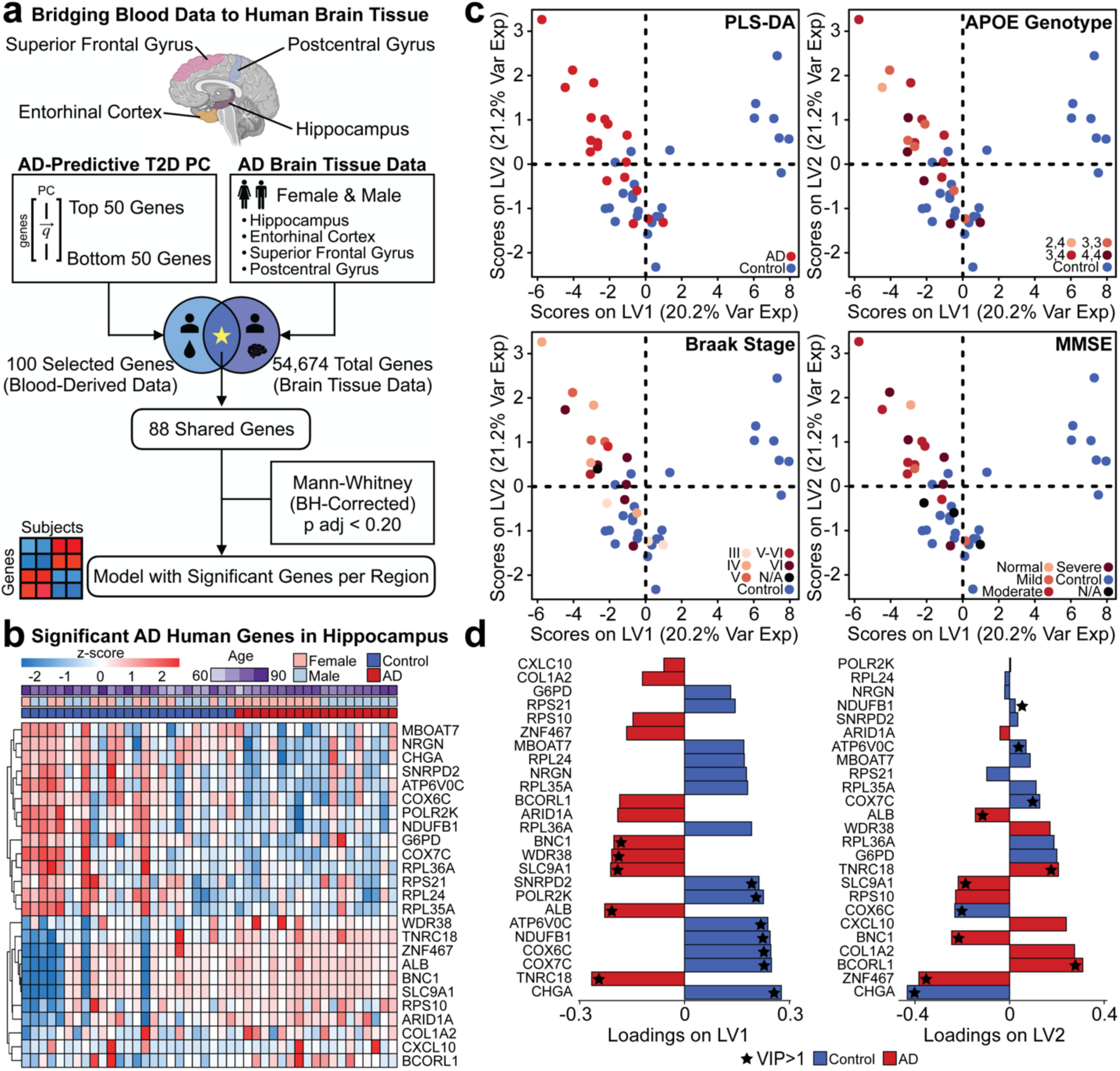
Translating blood-predictable signatures to the brain. **(a)** Method of testing blood-derived data predictability in the brain. **(b)** Z-score of significant AD-associated genes identified in the human hippocampal dataset (Mann-Whitney adjusted by Benjamini-Hochberg, *p* adjusted < 0.20). **(c)** PLS-DA model using significant genes to predict AD status. AD groups are labeled by APOE genotype, Braak stage, and MMSE. **(d)** Loading variables LV1 and LV2 for the model are presented. A VIP>1 is annotated with a star, and the color of the loading bar represents the highest contribution to the specific class by the respective gene.

We annotated the subjects within the PLS-DA plot by their respective apolipoprotein E (APOE) genotype, Braak stage, and mini-mental state examination (MMSE) scores (**Fig. 6d**). These were used since APOE e4 is the greatest genetic risk factor for AD^44^, Braak stage assesses neurofibrillary tangle pathology^45^, and MMSE for cognitive impairment screening^46^. There was clear separation between AD and control groups in our PLS-DA model and we identified a subset of genes loaded in the latent variables (LVs) most predictive of disease status (**Fig. 6d**). On LV1, we identified genes with variable importance of projection (VIP) greater than 1 associated with the control group, including *SNRPD2*, *POLR2K*, *ATP6V0C*, *NDUFB1*, *COX6C*, *COX7C*, and *CHGA*. For the AD group, we found *BNC1*, *WDR38*, *SLC9A1*, *ALB*, and *TNRC18* with a VIP>1. Although there was no separation across the disease classes on LV2, we found *NDUFB1*, *ATP6V0C*, *COX7C*, *COX6C*, and *CHGA* contributed greater than average (VIP > 1) to the control group, whereas *ALB*, *TNRC18*, *SLC9A1*, *BNC1*, *BCORL1*, and *ZNF467* had a VIP >1 for AD.

After observing separation across disease classes in the hippocampus brain data, we next determined if the T2D blood biomarkers able to stratify AD conditions in blood were reflective in other parts of the brain. We built PLS-DA models for the EC, SFG, and PoCG. Of the 88 genes that matched in the human brain tissue data, five genes were significant across AD and control groups in the EC (**Fig. 7a**). Using these genes for the PLS-DA model, we found distinct separation across LV1, and identified *RIN3*, *RPL36A*, and *POLR2K* as genes with a VIP greater than 1 (**Fig. 7b**). In the SFG brain region, we identified four significant genes: *RIN3*, *CSTA*, *RCN3*, and *RPL36A* (**Fig. 7c**). In the SFG model, *RIN3* and *RPL36A* contributed most to separation between the AD and control groups (**Fig. 7d**). In the PoCG region, three genes significantly separated AD and control, including *PRAM1*, *RCN3*, and *RPL36A* (**Fig. 7e-f**). For each of these three brain regions, additional annotation on the PLS-DA subjects by APOE genotype, Braak stage, and MMSE were visualized for the EC, SFG, and PoCG PLS-DA models (**Supplementary Fig. S5**).

**Figure 7.**
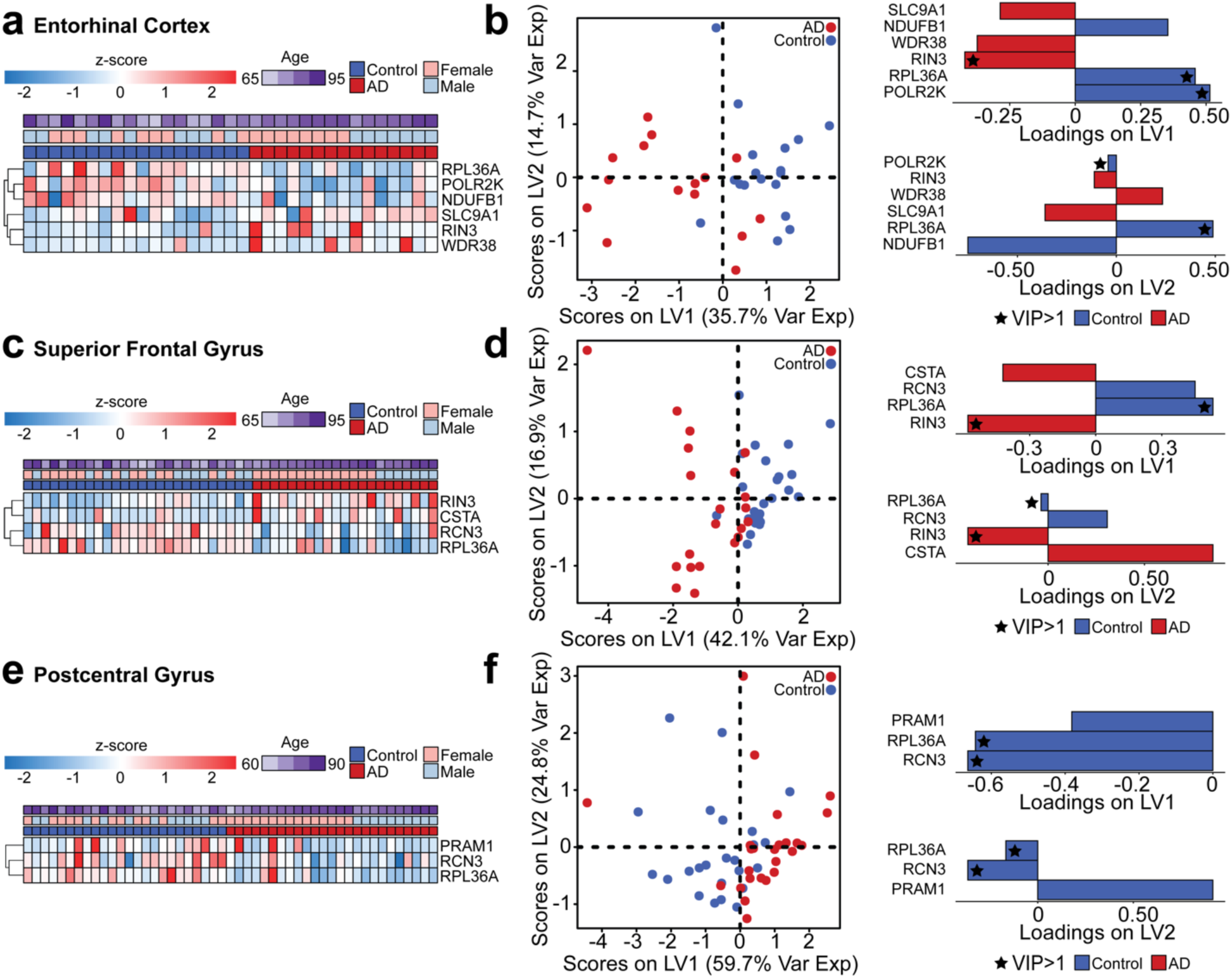
PLS-DA models using blood biomarkers to predict AD status in other brain regions. **(a)** Z-score of significant genes identified in the human EC dataset. **(b)** PLS-DA using the significant genes on the EC data with loadings on LV1 and LV2. **(c)** Z-score of significant genes identified in the human SFG dataset. **(d)** PLS-DA using the significant genes on the SFG data with loadings on LV1 and LV2. **(e)** Z-score of significant genes identified in the human PoCG dataset. **(f)** PLS-DA using the significant genes on the PoCG data with loadings on LV1 and LV2. For all brain regions, the significance of the genes was determined by a Mann-Whitney adjusted by Benjamini-Hochberg (*p* adjusted < 0.20) across AD and control groups.

## DISCUSSION

In this study, we used blood transcriptomics data from human T2D and AD studies to understand the potential pathways by which T2D affects AD pathology. Our cross-disease model identified a T2D-derived blood gene signature predictive of AD status and therapeutic candidates associated with non-T2D and AD status. A subset of genes in the T2D blood were predictive of AD status in four brain regions, showing the cross-disease model’s significance and implications.

Chemokine signaling pathways were involved in patients of T2D^47^ by routes of downstream inflammation^48^ and AD^49^ with connections to cognitive decline. Wnt signaling also played a role in metabolic dysregulation^50^ and loss of synaptic integrity^51^. Insulin pathways were enriched in AD conditions, consistent with prior literature showing insulin resistance^52^ is associated with an increased risk for AD development^53^. Pathways, such as MAPK and NOTCH, were enriched in AD conditions, with MAPK-p38 phosphorylation associated with both T2D and AD^54,55^. *Notch1* expression decreases beta cell masses and insulin secretion in rodents^56^ and was significantly different across control and AD groups in our analysis^57^. FC epsilon RI is also altered in T2D and AD cases, such that downstream mast cells are affected^58^.

We also identified cellular processes and metabolism pathways on the AD predictive T2D PC2. Elevated neutrophil activation to chemokines and transendothelial migration is associated with T2D^59^. In AD, monocytes and human brain microvascular endothelial cells expressing *CXCL1* are associated with amyloid-beta-induced migration from the blood to the brain^60^. FC gamma receptor-mediated phagocytosis is observed in T2D in compromised monocyte phagocytosis^61^. *PRKCD* is associated with amyloid-beta significantly triggered neurodegeneration in AD^62^. In blood, coagulation is active in hyperglycemia^63^ and factor XIII Val34Leu gene polymorphism is associated with sporadic AD^64^. Lastly, heme metabolism was associated with T2D and AD. A T2D-based study reported that increased dietary heme iron intake increased the risk of T2D^65^. In an AD study, altered heme metabolism was noted in AD brain samples^66^ (**Supplementary Table S6**).

From our drug screening analysis, we identified T2D and AD medications whose perturbed gene signatures significantly associated with the healthy state on the cross-disease predictive T2D PC2. The T2D (alogliptin and glipizide) and AD (galantamine and donepezil) medications that induced gene signatures correlated with T2D PC2 are current therapies for T2D and AD^67^. Alogliptin, an FDA-approved T2D, has been shown to reduce hippocampal insulin resistance in amyloid-beta-induced AD rodent models^68^. Glipizide has conflicting findings, with one study showed improved glycemic control and memory^69^ and another reported the drug be associated with higher risk of AD than metformin, another T2D medication^70^. Overall, the identification of these medications in our analysis shows promise for high-throughput drug screening integrated in a cross-disease modeling framework for comorbid conditions.

Our PLS-DA models identified signatures encoded in the T2D PC2 predicted AD status in brain tissue and many genes from our blood-based signature have associations with AD pathology in the brain. Individuals with MCI and AD show decreased *SNRPD2* expression levels in the hippocampus^71–73^, as well as decreased *POLR2K*^74,75^. *COX* deficiency has been reported in both AD brain and blood samples^76^. *CHGA* was associated with senile and pre-amyloid plaques^77^ and linked to AD compared to control groups in cerebrospinal fluid^78^. Our findings in literature show that *ALB* may differ across blood and brain^79,80^. While others reported decreased serum *ALB* levels increased the risk of AD, our findings in the hippocampus showed the opposite effects.

In the EC, SFG, and PoCG brain regions, *RIN3* was reported to have significantly elevated mRNA levels in the hippocampus and cortex of *APP/PS1* mouse models for AD^81^ and is a signature gene expressed in peripheral blood and the brain^81,82^. In a metformin response, drug-naïve T2D study, *RPL36A* correlated with a change in hemoglobin A1c levels^83^. In AD, *RPL36A* was found to be downregulated in cells stimulated by amyloid-beta^84^. This downregulation was consistent with our findings in the AD groups (**Supplementary Table S7**). These findings suggest that some gene signatures in T2D blood predictive of AD are present in the brain, linking blood-based biomarkers to primary tissue pathobiology.

A limitation to our study is that that data from large-scale human studies simultaneously studying the relationship between T2D and AD are still rare, meaning sample sizes and demographic representation of the human population across sex, age, and other variables is limited. Addressing this gap in the AD-T2D axis would improve opportunities to integrate other clinical variables, such as hemoglobin A1c for T2D, pathological results of amyloid-beta quantification for AD, and other human demographic variables known to be linked to AD and T2D pathology.

Our work introduced a new application for cross-disease modeling using TransComp-R to identify significantly relevant shared pathways by which T2D influences AD development. We found gene signatures in the peripheral blood of T2D subjects predictive of AD pathology, and identified a subset of genes in the blood that significantly predicted AD status in four brain regions. These findings shed insight into the shared comorbidity between T2D and AD and encourage future applications of TransComp-R for cross-disease modeling.

## MATERIALS AND METHODS

### Data selection

Human AD and T2D transcriptomic datasets were selected on GEO with the requirements that samples were collected from similar blood sample collection processes, a sample size of 10 or greater per condition, and demographic information containing sex and age. The datasets on GEO were scanned by using combinations of phrases, including “Alzheimer’s disease,” “diabetes,” “blood,” and “gene expression.” Like the blood data, post-mortem human brain tissue gene expression was identified using the information criteria containing human data with a cohort size greater than 10 per condition. Terms used to identify data on GEO included “brain,” “Alzheimer’s disease,” “human,” and “gene expression.”

### Pre-processing and normalization

Transcriptomic AD and T2D human data were acquired from GEO using Bioconductor tools in R (*GEOquery* ver. 2.70.0, *limma* ver. 3.58.1, and *Biobase* ver. 2.62.0)^85–87^. To reduce potential bias from younger age participants in the data, we removed all subjects 55 years old or below from the study in both the AD and T2D datasets with the justification of balancing the established age of late onset of AD (65 years). The T2D baseline group was used. For the AD cohorts, conditions that were not AD or control were excluded from the study. The datasets were then log_2_ transformed and matched for the same gene overlap. The genes shared across all AD and T2D datasets were normalized by z-score before computational modeling with TransComp-R.

### Cross-disease modeling with TransComp-R

We conducted TransComp-R by applying PCA on the T2D data with both disease and control groups. The number of PCs that encoded transcriptomic variation between healthy and T2D subjects was limited to a total explained cumulative variance of 80%. The two AD datasets were individually projected into the T2D PCA space, such that there were two separate models: T2D with AD cohort 1 and T2D with AD cohort 2. The projection of AD data into the T2D PCA space can be described by matrix multiplication:

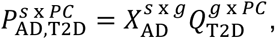

where matrix *P^s^* ^x^ *^PC^*, the projection of AD data onto the T2D space, defined by columns of T2D PCs and rows of AD subjects, is represented by the product of matrix *X^s^* ^x^ *^g^* and *Q^g^* ^x^ *^PC^*. Here, *s* is represented by AD subjects, *g* is represented by the gene list shared by AD and T2D, and *PC* is the principal components from the T2D space.

### Variance explained in Alzheimer’s disease by principal components of type 2 diabetes

To determine the translatability of T2D variance onto the AD data, we quantified the percent variability that is explained in AD by the T2D PCs with the following equation:

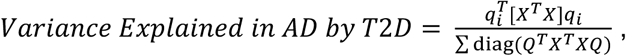

where AD data matrix *X*, projected onto a matrix *Q* containing columns of T2D PCs by matrix multiplication (*T* representing a matrix transpose). The percent variance of AD in *X* explained by a PC (*q_i_*) of *Q* was then calculated.

### Variable selection of T2D PCs

The T2D PCs predictive of AD outcomes were identified by employing LASSO across twenty random rounds of ten-fold cross-validations regressing the AD positions in T2D PC space against AD disease status. Demographic sex and age variables describing the subjects from the AD datasets were included in the GLM:

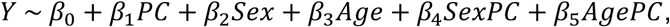

PCs with a coefficient frequency greater than 4 of the 20 rounds (25% selection frequency) in at least two of the three PC terms (PC, Sex*PC, or Age*PC) were selected for GLMs with individual PCs regressed against AD outcomes. T2D PCs that were consistently significant in both AD cohorts (*p* value < 0.05) were selected for further biological interpretation.

### Gene set enrichment analysis

Loadings of the PCs selected by the GLM were analyzed with GSEA in R (*msigdbr* ver. 7.5.1, *fgsea* ver. 1.28.0, and *clusterProfiler* ver. 4.10.1)^88–90^. Two data collections (KEGG and Hallmark) were downloaded from the Molecular Signatures Database to identify enriched biological pathways. Identified pathways were determined to be significant, with a Benjamini-Hochberg adjusted *p* value of less than 0.01 to account for multiple hypothesis testing. The imputed parameters to run GSEA included a minimum gene size of 5, a maximum gene size of 500, and epsilon, the tuning constant of 0. The default setting of 1000 permutations was used.

### Identifying shared genes across enriched biological pathways

We used *igraph* (ver. 2.0.3)^91^ in R to identify overlapping genes that may be commonly enriched across multiple biological pathways identified from GSEA. We then processed the R-generated data in *Cytoscape* (ver. 3.10.2)^92^ to enhance pathway visualization. We established the nodes representing different biological pathways and the edge thickness by the number of overlapping genes between the two biological pathways. Additionally, the node size was determined by the number of total enriched genes contributing to the biological pathway as determined by GSEA, with the node colors red and blue used to discern pathway associations with AD or control groups, respectively.

### Fold-change comparison cross-disease

The relationship of different gene expression across AD and T2D conditions was compared using the log_2_ fold change of each gene shared across the AD and T2D blood data. For each dataset (T2D and AD), the log_2_ fold change of each gene expression was calculated by taking the log_2_ of the average gene expression of the disease groups divided by the average gene expression of the control groups. Different gene expression relationships were compared across the T2D and AD datasets.

### Sex-based comparison across type 2 diabetes principal component scores

PC scores were compared across sex and disease conditions to compare PC predictability across sex demographics. A Mann-Whitney pair-wise test was used to compare AD females to control females and AD males to control males. To account for multiple hypothesis testing, a Benjamini-Hochberg adjusted *p* value less than 0.05 was determined significant for the analysis.

### Computational gene expression correlational analysis

Potentially therapeutic drugs correlated with T2D PCs predictive of AD were screened using publicly available data from the L1000 Consensus Signatures Coefficient Tables (Level 5) from the LINCS database. Before screening, the LINCS drug data was pre-processed by excluding all drugs with no known targets based on the LINCS small molecules metadata.

To identify candidate drugs associated with T2D and AD, two data sources were compiled: DEGs from each respective drug from LINCS and the loadings from the T2D PCs predictive of AD. DEGs for each drug were determined through the following: The characteristic direction values, which signified the drug’s up- or down-regulation of a gene, were scaled to obtain their z-score values^40^. The list of DEGs for each drug was then identified if the gene’s z-score value presented with a *p* value less than 0.05. The original characteristic direction values for the selected genes for each respective drug were then isolated. For each T2D PC that was able to stratify transcriptomic variance between control and AD subjects, differentially expressed drug genes and PC gene loadings were matched. A Spearman correlation was calculated to determine the correlation between PC loadings and the DEGs’ characteristic direction coefficients for each drug. For a given T2D PC of interest, drugs were ranked by their respective Spearman’s ρ values. The correlations’ *p* values were corrected by Benjamini-Hochberg before visualizing the drugs’ ranks against their 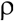 values (adjusted *p* value < 0.05).

### Filtering genetic blood biomarkers for computational modeling of brain tissue data

The top 50 and bottom 50 genes, ranked by their respective scores on the T2D PC predictive of AD in blood, were used to filter genes of AD brain tissue data. After filtering for matching genes, a Benjamini-Hochberg adjusted Mann-Whitney test was performed to determine significant genes.

An adjusted *p* value of less than 0.20 was deemed significant to allow for a more permissible list of potential genes that relate the blood to the brain. The significant genes were then used for PLS-DA modeling.

### Partial least squares discriminant analysis

Using R (*mixOmics* ver. 6.26.0)^93^, we constructed a PLS-DA model to determine the predictability of blood-based gene expression markers in the human brain. Specifically, we used PCs derived from T2D blood transcriptomic data predictive of AD outcomes in blood profiles and selected the top 50 and bottom 50 gene loadings as a filter for hippocampal tissue transcriptomic data in human subjects. A PLS-DA model screening for the 100 genes was used to determine if all genes driving the transcriptomic variation in the T2D PC could stratify AD and control in brain tissue. As an additional follow-up, the 100 filtered genes selected by the blood data significantly distinguishable among AD and control in human blood were also used to construct the PLS-DA model. The number of latent variables used for the model was determined by 100 randomly repeated three-fold cross-validation based on the model with the lowest cross-validation error rate.

As a way to determine the most important predictors driving separation and predictive accuracy in the PLS-DA model, we calculated the VIP score for each gene. For a given number of PLS-DA components *A*, the VIP for each gene predictor, *k*, is calculated by:

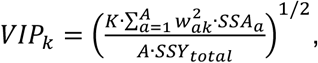

where *K* is the total number of gene predictors, *w_ak_* is the weight of predictor *k* in the *a^th^* LV component. The total sum of squares explained in all LV components is represented by *SSY_total_*. A calculated VIP score greater than 1 signifies that a given gene is an important variable for a specific LV in the PLS-DA model.

AD subjects were annotated by their APOE genotype, Braak stage, and MMSE score among each PLS-DA model. The MMSE numerical scores, which evaluate cognitive impairment, were aggregated based on standardized scoring metrics such that 30-26 was normal, 25-20 was mild, 19-10 was moderate, and 9-0 was severe^94^. The control groups did not have any clinical records.

## ABBREVIATIONS

AD: Alzheimer’s disease
APOE: Apolipoprotein E
BBB: Blood-brain barrier
DEG: Differentially expressed gene
EC: Entorhinal cortex
FDA: Food and Drug Administration
GEO: Gene Expression Omnibus
GLM: Generalized linear model
GSEA: Gene set enrichment analysis
LASSO: Least Absolute Shrinkage and Selection Operator
LINCS: Library of Integrated Network-Based Cellular Signatures
LV: Latent variable
MCI: Mild cognitive impairment
MMSE: Mini-Mental State Examination
PC: Principal component
PCA: Principal component analysis
PLS-DA: Partial least squares discriminant analysis
PoCG: Postcentral gyrus
SFG: Superior frontal gyrus
T2D: Type 2 diabetes
TransComp-R: Translatable Components Regression
VIP: Variable importance of projection

## ACKNOWLEDGEMENTS

This work is supported by an award from the Good Ventures Foundation and Open Philanthropy (DKB, BKB, and JHP). This work is also supported by R01AG072513 from the National Institute on Aging (EAP). BKB acknowledges the National Science Foundation for support under the Graduate Research Fellowship Program (GRFP) under grant number DGE-1842166. BKB also acknowledges the support of the NIH T32 predoctoral fellowship T32DK101001 from the National Institute of Diabetes and Digestive and Kidney Diseases. Additionally, BKB acknowledges a research grant award from the Research Hub Foundation.

## AUTHOR CONTRIBUTIONS

BKB: Conceptualization, data curation, formal analysis, funding acquisition, investigation, methodology, visualization, writing-original draft, writing-review & editing. JHP: Data curation, methodology, writing-review & editing. JHP: Conceptualization, data curation, methodology, writing-review & editing. EAP: Conceptualization, funding acquisition, methodology, project administration, resources, writing-review & editing. DKB: Conceptualization, funding acquisition, methodology, project administration, resources, writing-review & editing.

## DATA AVAILABILITY

We accessed all blood-derived T2D RNA-seq and AD microarray expression data from Gene Expression Omnibus under accession numbers GSE184050, GSE63060, and GSE63061. Additionally, hippocampal human data was acquired from Gene Expression Omnibus with the accession number GSE48350. The computational correlational analysis data was acquired from the Library of Integrated Network-Based Cellular Signatures database’s L1000 Consensus Signatures Coefficient Tables (Level 5).

## CODE AVAILABILITY

All code used for analysis is made publicly available at https://github.com/Brubaker-Lab/CrossDisease-TransCompR-T2D-AD-Human.

## SUPPLEMENTARY INFORMATION

**Supplementary Figure S1.**
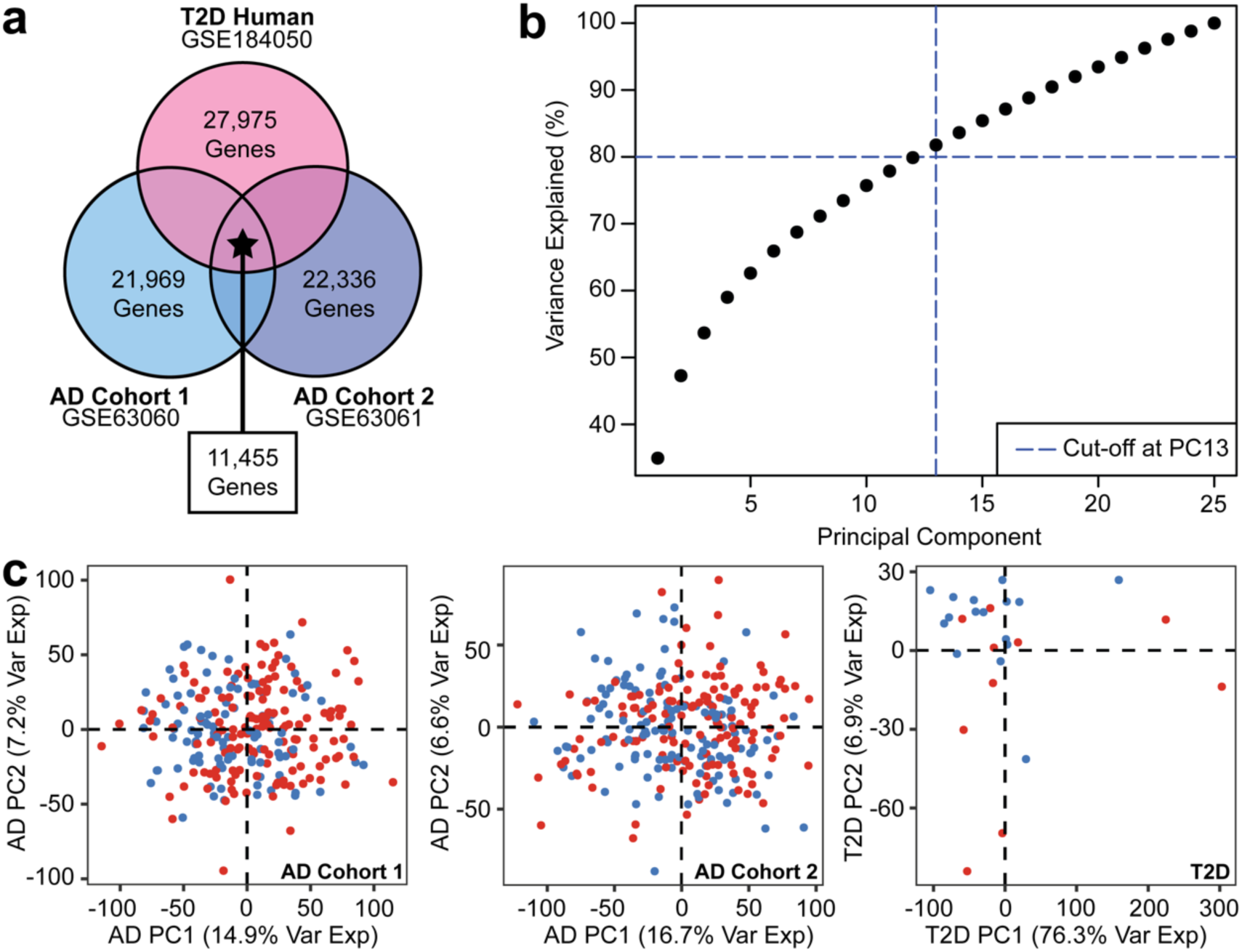
Data processing of the AD and T2D data. **(a)** Gene overlaps across the three publicly available transcriptomics data. **(b)** Cumulative variance was explained for T2D PCs with a threshold of 80%. **(c)** PCA of the AD cohort 1, AD cohort 2, and T2D data.

**Supplementary Figure S2.**
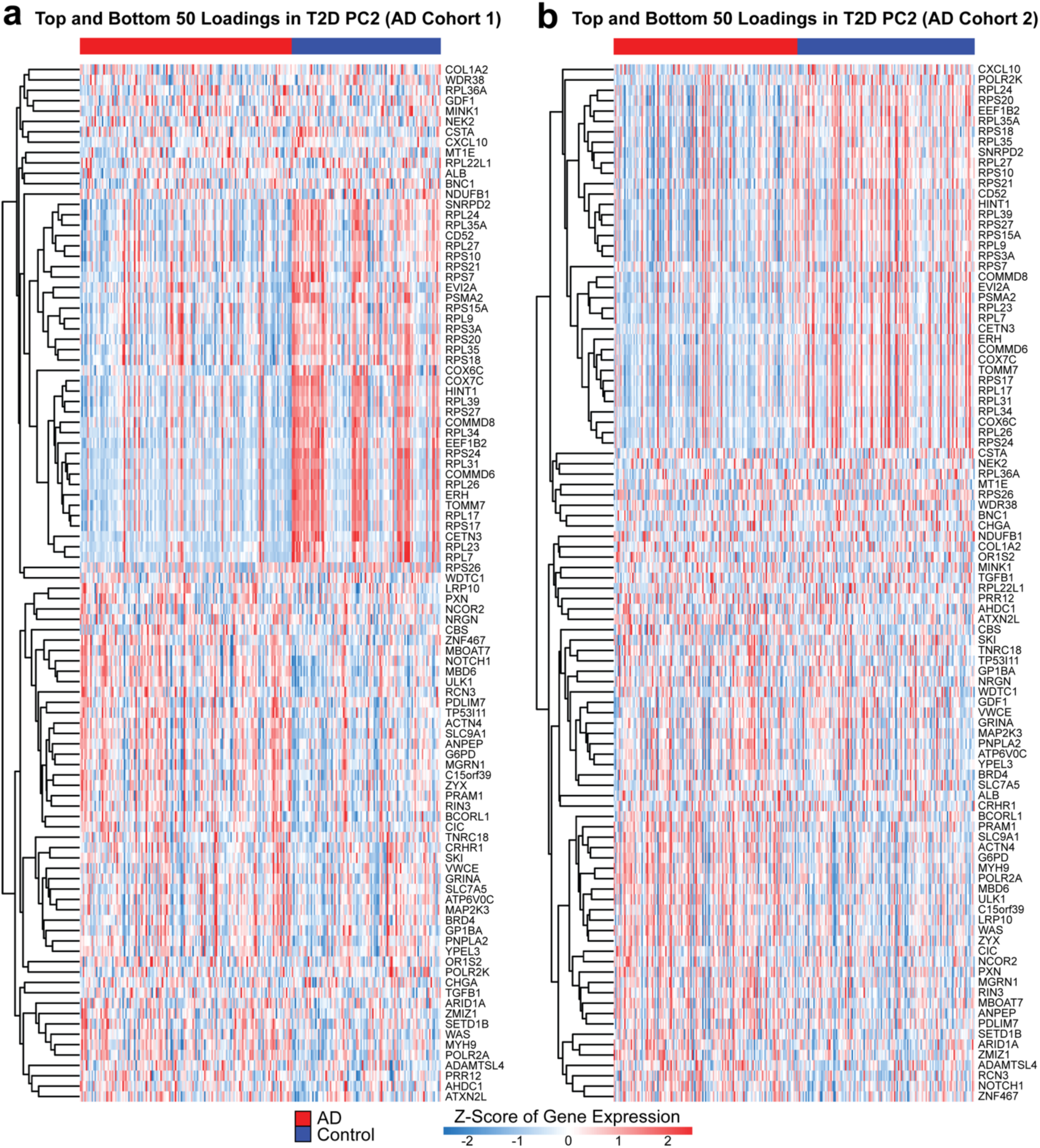
Identification of genes separating between healthy and AD groups. Hierarchical clustering of the top and bottom 50 T2D PC2 loadings in T2D PC2 for **(a)** AD cohort 1 and **(b)** AD cohort 2.

**Supplementary Figure S3.**
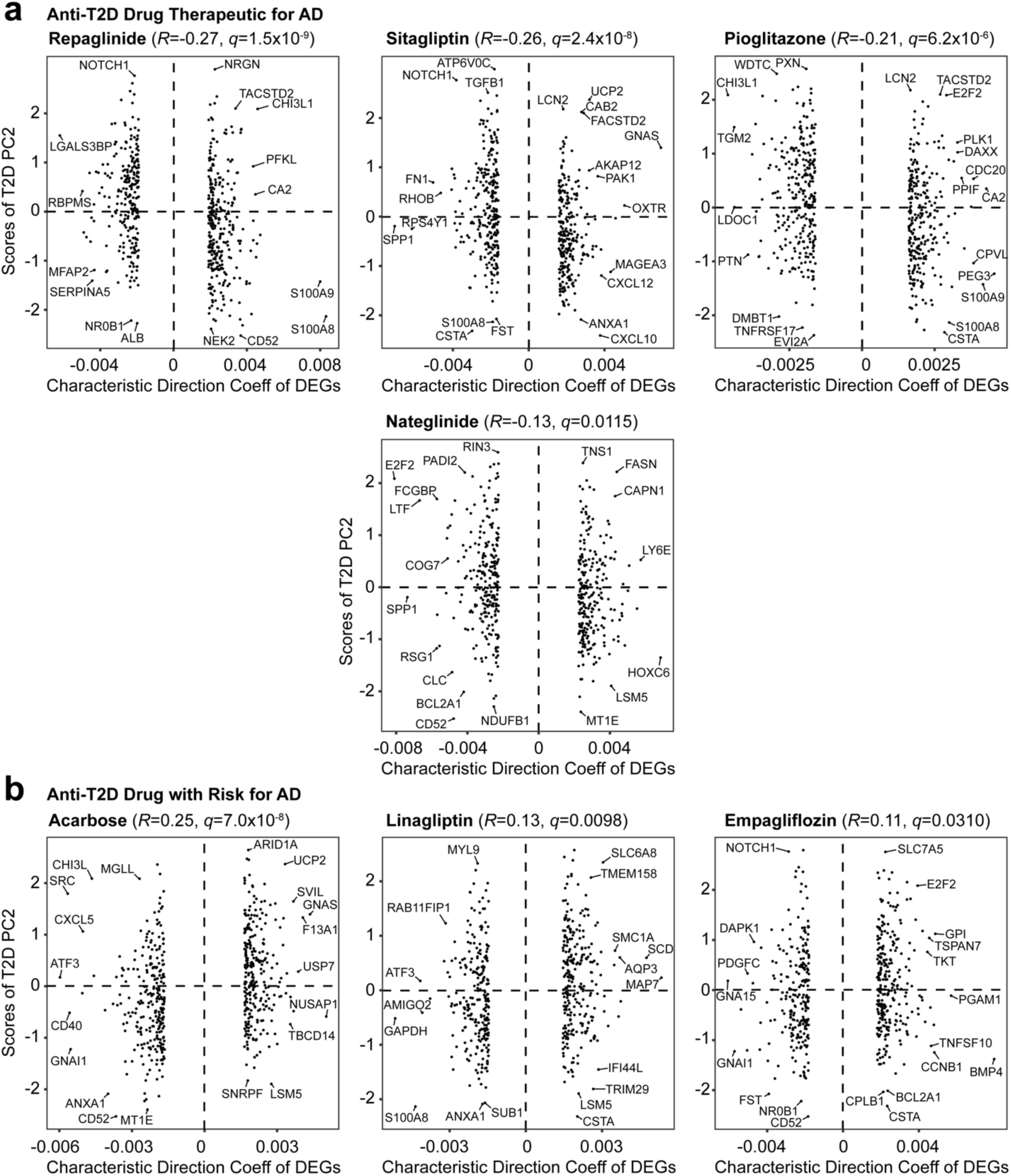
Additional anti-T2D drugs approved by the FDA. **(a)** Anti-T2D therapeutics with potential reduction of AD pathology. **(b)** Anti-T2D therapeutics with increased risk for AD.

**Supplementary Figure S4.**
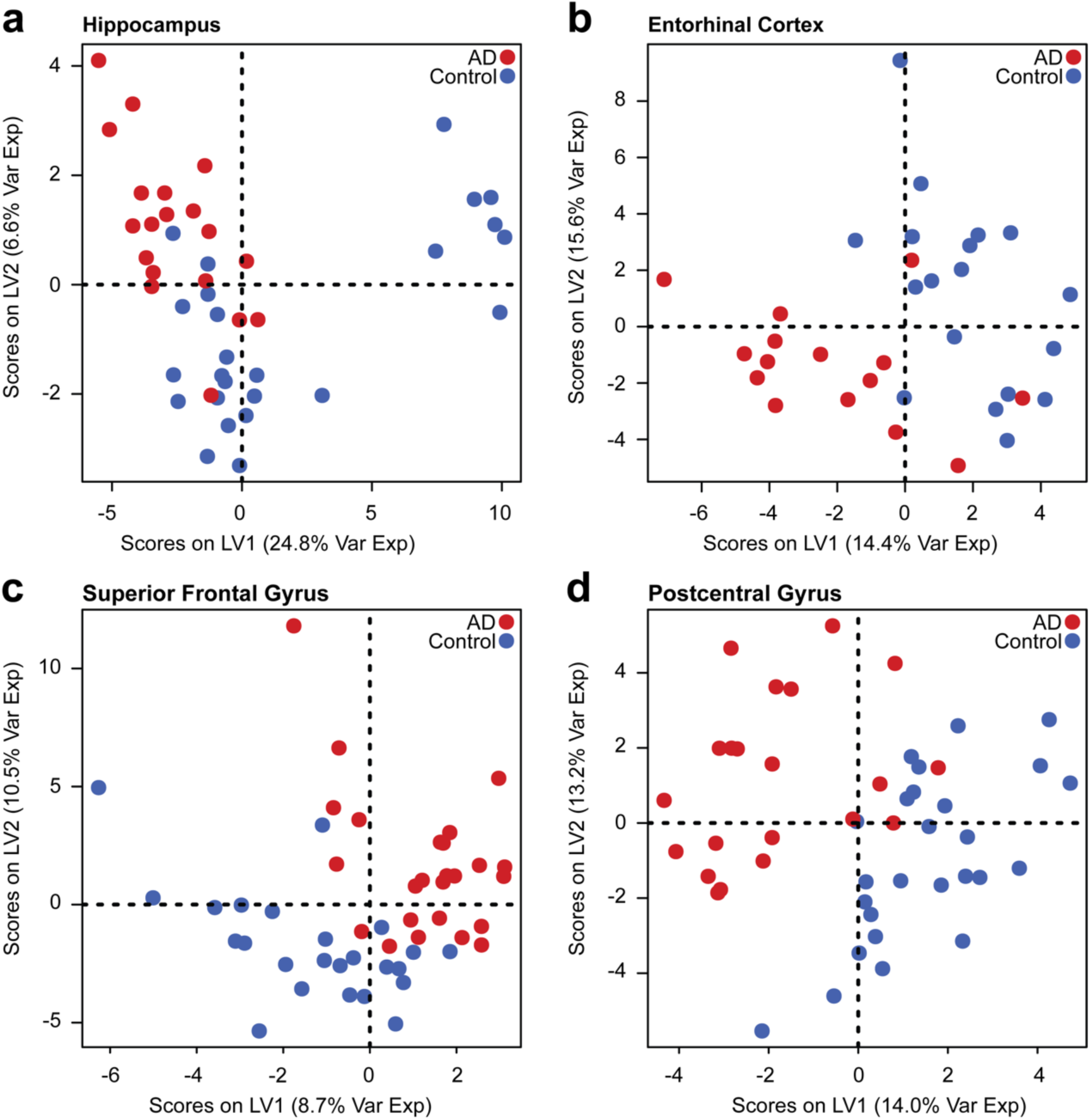
PLS-DA models of the brain tissue gene expression filtered by the 88 represented in the T2D PC2 loading. Models constructed for (a) the hippocampus, (b) the entorhinal cortex, (c) the superior frontal gyrus, and (d) the postcentral gyrus.

**Supplementary Figure S5.**
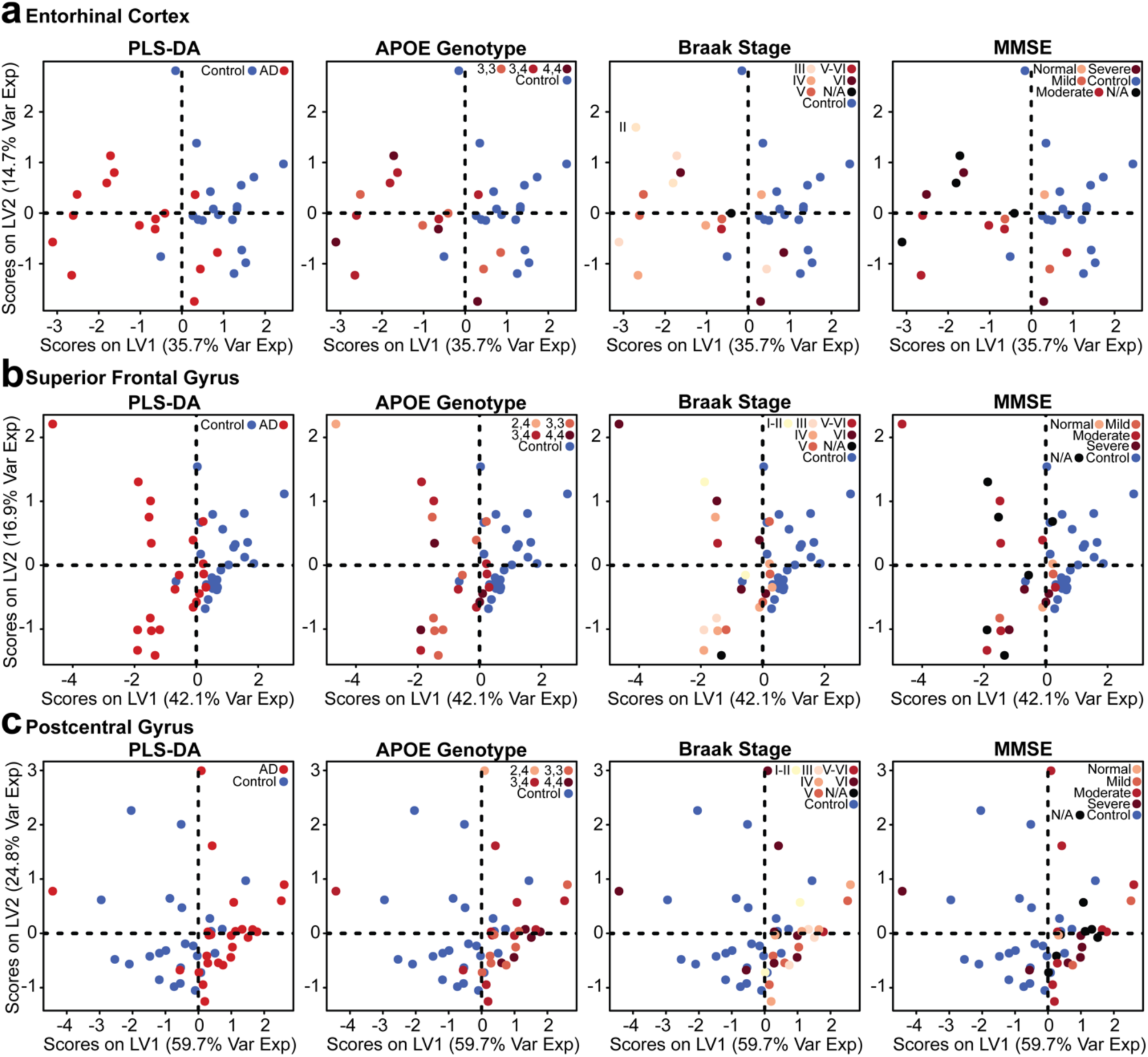
Annotated PLS-DA models by genotype and clinical scores. Subjects are further labeled by their respective APOE genotype, Braak stage, and MMSE for brain tissue collected for the **(a)** entorhinal cortex, **(b)** superior frontal gyrus, and **(c)** postcentral gyrus.

## REFERENCES

1. Wang, K.-C. et al. Risk of Alzheimer’s Disease in Relation to Diabetes: A Population-Based Cohort Study. Neuroepidemiology 38, 237–244 (2012).

2. Janson, J. et al. Increased Risk of Type 2 Diabetes in Alzheimer Disease. Diabetes 53, 474– 481 (2004).

3. Gudala, K., Bansal, D., Schifano, F. & Bhansali, A. Diabetes mellitus and risk of dementia: A meta-analysis of prospective observational studies. Journal of Diabetes Investigation 4, 640– 650 (2013).

4. Barbiellini Amidei, C., et al. Association Between Age at Diabetes Onset and Subsequent Risk of Dementia. JAMA 325, 1640–1649 (2021).

5. Cheng, G., Huang, C., Deng, H. & Wang, H. Diabetes as a risk factor for dementia and mild cognitive impairment: a meta-analysis of longitudinal studies. Internal Medicine Journal 42, 484–491 (2012).

6. Teixeira, M. M. et al. Association between diabetes and cognitive function at baseline in the Brazilian Longitudinal Study of Adult Health (ELSA-Brasil). Sci Rep 10, 1596 (2020).

7. Sun, D. et al. Type 2 Diabetes and Hypertension: A Study on Bidirectional Causality. Circulation Research 124, 930–937 (2019).

8. Srodulski, S. et al. Neuroinflammation and neurologic deficits in diabetes linked to brain accumulation of amylin. Molecular Neurodegeneration 9, 30 (2014).

9. Palazzuoli, A. & Iacoviello, M. Diabetes leading to heart failure and heart failure leading to diabetes: epidemiological and clinical evidence. Heart Fail Rev 28, 585–596 (2023).

10. Chen, R., Ovbiagele, B. & Feng, W. Diabetes and Stroke: Epidemiology, Pathophysiology, Pharmaceuticals and Outcomes. Am J Med Sci 351, 380–386 (2016).

11. Kumar, M. et al. The Bidirectional Link Between Diabetes and Kidney Disease: Mechanisms and Management. Cureus 15, e45615.

12. Riching, A. S., Major, J. L., Londono, P. & Bagchi, R. A. The Brain–Heart Axis: Alzheimer’s, Diabetes, and Hypertension. ACS Pharmacol Transl Sci 3, 21–28 (2019).

13. Candeias, E. et al. The impairment of insulin signaling in Alzheimer’s disease. IUBMB Life 64, 951–957 (2012).

14. De Felice, F. G., Gonçalves, R. A. & Ferreira, S. T. Impaired insulin signalling and allostatic load in Alzheimer disease. Nat Rev Neurosci 23, 215–230 (2022).

15. Morgen, K. & Frölich, L. The metabolism hypothesis of Alzheimer’s disease: from the concept of central insulin resistance and associated consequences to insulin therapy. J Neural Transm 122, 499–504 (2015).

16. Pontzer, H. et al. Daily energy expenditure through the human life course. Science 373, 808– 812 (2021).

17. Liu, P. et al. High-fat diet-induced diabetes couples to Alzheimer’s disease through inflammation-activated C/EBPβ/AEP pathway. Mol Psychiatry 27, 3396–3409 (2022).

18. De Sousa, R. A. L. et al. An update on potential links between type 2 diabetes mellitus and Alzheimer’s disease. Mol Biol Rep 47, 6347–6356 (2020).

19. Khan, M. S. H. & Hegde, V. Obesity and Diabetes Mediated Chronic Inflammation: A Potential Biomarker in Alzheimer’s Disease. Journal of Personalized Medicine 10, (2020).

20. Bury, J. J. et al. Type 2 diabetes mellitus-associated transcriptome alterations in cortical neurones and associated neurovascular unit cells in the ageing brain. Acta Neuropathol Commun 9, 5 (2021).

21. Rom, S. et al. Hyperglycemia-driven neuroinflammation compromises BBB leading to memory loss in both diabetes mellitus (DM) type 1 and type 2 mouse models. Mol Neurobiol 56, 1883–1896 (2019).

22. Liu, Y., Huber, C. C. & Wang, H. Disrupted blood-brain barrier in 5×FAD mouse model of Alzheimer’s disease can be mimicked and repaired *in vitro* with neural stem cell-derived exosomes. Biochemical and Biophysical Research Communications 525, 192–196 (2020).

23. Blanchette, M. & Daneman, R. Formation and maintenance of the BBB. Mechanisms of Development 138, 8–16 (2015).

24. Kadry, H., Noorani, B. & Cucullo, L. A blood–brain barrier overview on structure, function, impairment, and biomarkers of integrity. Fluids and Barriers of the CNS 17, 69 (2020).

25. Janelidze, S. et al. Increased blood-brain barrier permeability is associated with dementia and diabetes but not amyloid pathology or APOE genotype. Neurobiology of Aging 51, 104–112 (2017).

26. Hu, Y., Zheng, Y., Wang, T., Jiao, L. & Luo, Y. VEGF, a Key Factor for Blood Brain Barrier Injury After Cerebral Ischemic Stroke. Aging Dis 13, 647–654 (2022).

27. Tousoulis, D. et al. Diabetes Mellitus-Associated Vascular Impairment: Novel Circulating Biomarkers and Therapeutic Approaches. Journal of the American College of Cardiology 62, 667–676 (2013).

28. Tryggestad, J. B. et al. Circulating adhesion molecules and associations with HbA1c, hypertension, nephropathy, and retinopathy in the Treatment Options for type 2 Diabetes in Adolescent and Youth study. Pediatric Diabetes 21, 923–931 (2020).

29. Hanlon, P. et al. Representation of people with comorbidity and multimorbidity in clinical trials of novel drug therapies: an individual-level participant data analysis. BMC Medicine 17, (2019).

30. Zilkens, R. R., Davis, W. A., Spilsbury, K., Semmens, J. B. & Bruce, D. G. Earlier Age of Dementia Onset and Shorter Survival Times in Dementia Patients With Diabetes. American Journal of Epidemiology 177, 1246–1254 (2013).

31. Huang, C., Luo, J., Wen, X. & Li, K. Linking Diabetes Mellitus with Alzheimer’s Disease: Bioinformatics Analysis for the Potential Pathways and Characteristic Genes. Biochem Genet 60, 1049–1075 (2022).

32. Karki, R. et al. Data-Driven Modeling of Knowledge Assemblies in Understanding Comorbidity Between Type 2 Diabetes Mellitus and Alzheimer’s Disease. Journal of Alzheimer’s Disease 78, 87–95 (2020).

33. Lee, T. & Lee, H. Shared Blood Transcriptomic Signatures between Alzheimer’s Disease and Diabetes Mellitus. Biomedicines 9, 34 (2021).

34. Brubaker, Douglas. K., et al. An interspecies translation model implicates integrin signaling in infliximab-resistant inflammatory bowel disease. Sci Signal 13, eaay3258 (2020).

35. Lee, M. J. et al. Computational Interspecies Translation Between Alzheimer’s Disease Mouse Models and Human Subjects Identifies Innate Immune Complement, TYROBP, and TAM Receptor Agonist Signatures, Distinct From Influences of Aging. Frontiers in Neuroscience 15, (2021).

36. Suarez-Lopez, L. et al. Cross-species transcriptomic signatures predict response to MK2 inhibition in mouse models of chronic inflammation. iScience 24, 103406 (2021).

37. Ball, B. K., Proctor, E. A. & Brubaker, D. K. Cross-Species Modeling Identifies Gene Signatures in Type 2 Diabetes Mouse Models Predictive of Inflammatory and Estrogen Signaling Pathways Associated with Alzheimer’s Disease Outcomes in Humans. in Biocomputing 2025 426–440 (WORLD SCIENTIFIC, 2024). doi:10.1142/9789819807024_0031.

38. Chen, H.-H. et al. Novel diabetes gene discovery through comprehensive characterization and integrative analysis of longitudinal gene expression changes. Human Molecular Genetics 31, 3191 (2022).

39. Sood, S. et al. A novel multi-tissue RNA diagnostic of healthy ageing relates to cognitive health status. Genome Biology 16, (2015).

40. 40. Xie, Z., et al. Getting Started with LINCS Datasets and Tools. Curr Protoc 2, e487 (2022).

41. Food and Drug Administration. Approved Drug Products with Therapeutic Equivalence Evaluations. (2024).

42. Berchtold, N. C. et al. Synaptic genes are extensively downregulated across multiple brain regions in normal human aging and Alzheimer’s disease. Neurobiol Aging 34, 1653–1661 (2013).

43. Cribbs, D. H. et al. Extensive innate immune gene activation accompanies brain aging, increasing vulnerability to cognitive decline and neurodegeneration: a microarray study. J Neuroinflammation 9, 179 (2012).

44. Jansen, W. J. et al. Prevalence of Cerebral Amyloid Pathology in Persons Without Dementia: A Meta-analysis. JAMA 313, 1924–1938 (2015).

45. Malek-Ahmadi, M., Perez, S. E., Chen, K. Mufson, E. J. Braak Stage, Cerebral Amyloid Angiopathy, and Cognitive Decline in Early Alzheimer’s Disease. J Alzheimers Dis 74, 189– 197 (2020).

46. Arevalo-Rodriguez, I. et al. Mini-Mental State Examination (MMSE) for the early detection of dementia in people with mild cognitive impairment (MCI). Cochrane Database Syst Rev 2021, CD010783 (2021).

47. Cereijo, R. et al. The chemokine CXCL14 is negatively associated with obesity and concomitant type-2 diabetes in humans. Int J Obes 45, 706–710 (2021).

48. Ball, B. K., Kuhn, M. K., Fleeman Bechtel, R. M., Proctor, E. A. & Brubaker, D. K. Differential responses of primary neuron-secreted MCP-1 and IL-9 to type 2 diabetes and Alzheimer’s disease-associated metabolites. Sci Rep 14, 12743 (2024).

49. Zhou, F., Sun, Y., Xie, X. & Zhao, Y. Blood and CSF chemokines in Alzheimer’s disease and mild cognitive impairment: a systematic review and meta-analysis. Alz Res Therapy 15, 107 (2023).

50. Fuster, J. J. et al. Noncanonical Wnt Signaling Promotes Obesity-Induced Adipose Tissue Inflammation and Metabolic Dysfunction Independent of Adipose Tissue Expansion. Diabetes 64, 1235–1248 (2014).

51. Liu, C.-C. et al. Deficiency in LRP6-Mediated Wnt Signaling Contributes to Synaptic Abnormalities and Amyloid Pathology in Alzheimer’s Disease. Neuron 84, 63–77 (2014).

52. Fischer, S. et al. Insulin-resistant patients with type 2 diabetes mellitus have higher serum leptin levels independently of body fat mass. Acta Diabetol 39, 105–110 (2002).

53. Schrijvers, E. M. C. et al. Insulin metabolism and the risk of Alzheimer disease. Neurology 75, 1982–1987 (2010).

54. Brown, A. E. et al. p38 MAPK activation upregulates proinflammatory pathways in skeletal muscle cells from insulin-resistant type 2 diabetic patients. American Journal of Physiology-Endocrinology and Metabolism 308, E63–E70 (2015).

55. Wang, S. et al. Peripheral expression of MAPK pathways in Alzheimer’s and Parkinson’s diseases. Journal of Clinical Neuroscience 21, 810–814 (2014).

56. Eom, Y. S. et al. Notch1 Has an Important Role in β-Cell Mass Determination and Development of Diabetes. Diabetes Metab J 45, 86–96 (2021).

57. Cho, S.-J. et al. Altered expression of Notch1 in Alzheimer’s disease. PLOS ONE 14, e0224941 (2019).

58. Kettner, A., Di Matteo, M. & Santoni, A. Insulin potentiates FcɛRI-mediated signaling in mouse bone marrow-derived mast cells. Molecular Immunology 47, 1039–1046 (2010).

59. Lin, Q. et al. Abnormal Peripheral Neutrophil Transcriptome in Newly Diagnosed Type 2 Diabetes Patients. Journal of Diabetes Research 2020, 9519072 (2020).

60. Zhang, K. et al. CXCL1 Contributes to β-Amyloid-Induced Transendothelial Migration of Monocytes in Alzheimer’s Disease. PLOS ONE 8, e72744 (2013).

61. Restrepo, B. I., Twahirwa, M., Rahbar, M. H. & Schlesinger, L. S. Phagocytosis via Complement or Fc-Gamma Receptors Is Compromised in Monocytes from Type 2 Diabetes Patients with Chronic Hyperglycemia. PLoS One 9, e92977 (2014).

62. Park, Y. H. et al. Dysregulated Fc gamma receptor–mediated phagocytosis pathway in Alzheimer’s disease: network-based gene expression analysis. Neurobiology of Aging 88, 24–32 (2020).

63. Stegenga, M. E. et al. Hyperglycemia Stimulates Coagulation, Whereas Hyperinsulinemia Impairs Fibrinolysis in Healthy Humans. Diabetes 55, 1807–1812 (2006).

64. Gerardino, L. et al. Coagulation factor XIII Val34Leu gene polymorphism and Alzheimer’s disease. Neurological Research 28, 807–809 (2006).

65. Wang, F. et al. Integration of epidemiological and blood biomarker analysis links haem iron intake to increased type 2 diabetes risk. Nat Metab 1–12 (2024) doi:10.1038/s42255-024-01109-5.

66. Atamna, H. & Frey, W. H. A role for heme in Alzheimer’s disease: Heme binds amyloid β and has altered metabolism. Proceedings of the National Academy of Sciences 101, 11153–11158 (2004).

67. Stanciu, G. D. et al. Link between Diabetes and Alzheimer’s Disease Due to the Shared Amyloid Aggregation and Deposition Involving Both Neurodegenerative Changes and Neurovascular Damages. J Clin Med 9, 1713 (2020).

68. Rahman, S. O. et al. Alogliptin reversed hippocampal insulin resistance in an amyloid-beta fibrils induced animal model of Alzheimer’s disease. European Journal of Pharmacology 889, 173522 (2020).

69. Gradman, T. J., Laws, A., Thompson, L. W. & Reaven, G. M. Verbal Learning and/or Memory Improves with Glycemic Control in Older Subjects with Non-Insulin-Dependent Diabetes Mellitus. Journal of the American Geriatrics Society 41, 1305–1312 (1993).

70. Akimoto, H. et al. Antidiabetic Drugs for the Risk of Alzheimer Disease in Patients With Type 2 DM Using FAERS. Am J Alzheimers Dis Other Demen 35, 1533317519899546 (2020).

71. Liu, N. et al. Hippocampal transcriptome-wide association study and neurobiological pathway analysis for Alzheimer’s disease. PLOS Genetics 17, e1009363 (2021).

72. Tao, Y. et al. The Predicted Key Molecules, Functions, and Pathways That Bridge Mild Cognitive Impairment (MCI) and Alzheimer’s Disease (AD). Front. Neurol. 11, (2020).

73. Bellou, E. & Escott-Price, V. Are Alzheimer’s and coronary artery diseases genetically related to longevity? Front. Psychiatry 13, (2023).

74. Wong, J. Altered Expression of RNA Splicing Proteins in Alzheimer’s Disease Patients: Evidence from Two Microarray Studies. DEE 3, 74–85 (2013).

75. Chaparro, R. J. et al. Nonobese diabetic mice express aspects of both type 1 and type 2 diabetes. Proc Natl Acad Sci U S A 103, 12475–12480 (2006).

76. Cardoso, S. M., Proença, M. T., Santos, S., Santana, I. & Oliveira, C. R. Cytochrome c oxidase is decreased in Alzheimer’s disease platelets. Neurobiology of Aging 25, 105–110 (2004).

77. Rangon, C.-M. et al. Different chromogranin immunoreactivity between prion and a-beta amyloid plaque. NeuroReport 14, 755 (2003).

78. Hölttä, M. et al. An Integrated Workflow for Multiplex CSF Proteomics and Peptidomics— Identification of Candidate Cerebrospinal Fluid Biomarkers of Alzheimer’s Disease. J. Proteome Res. 14, 654–663 (2015).

79. Kim, J. W. et al. Serum albumin and beta-amyloid deposition in the human brain. Neurology 95, e815–e826 (2020).

80. Min, J.-Y. et al. Chronic Status of Serum Albumin and Cognitive Function: A Retrospective Cohort Study. J Clin Med 11, 822 (2022).

81. Shen, R. et al. Upregulation of RIN3 induces endosomal dysfunction in Alzheimer’s disease. Transl Neurodegener 9, 26 (2020).

82. Kajiho, H. et al. RIN3: a novel Rab5 GEF interacting with amphiphysin II involved in the early endocytic pathway. Journal of Cell Science 116, 4159–4168 (2003).

83. Vohra, M. et al. Implications of genetic variations, differential gene expression, and allele-specific expression on metformin response in drug-naïve type 2 diabetes. Journal of Endocrinological Investigation 46, 1205 (2023).

84. Deng, L. et al. Amyloid β Induces Early Changes in the Ribosomal Machinery, Cytoskeletal Organization and Oxidative Phosphorylation in Retinal Photoreceptor Cells. Front Mol Neurosci 12, 24 (2019).

85. Davis, S. & Meltzer, P. S. GEOquery: a bridge between the Gene Expression Omnibus (GEO) and BioConductor. Bioinformatics 23, 1846–1847 (2007).

86. Ritchie, M. E. et al. limma powers differential expression analyses for RNA-sequencing and microarray studies. Nucleic Acids Research 43, e47 (2015).

87. Huber, W. et al. Orchestrating high-throughput genomic analysis with Bioconductor. Nat Methods 12, 115–121 (2015).

88. Dolgalev, I. msigdbr: MSigDB Gene Sets for Multiple Organisms in a Tidy Data Format. (2022).

89. Korotkevich, G. et al. Fast gene set enrichment analysis. 060012 Preprint at 10.1101/060012 (2021).

90. Yu, G., Wang, L.-G., Han, Y. & He, Q.-Y. clusterProfiler: an R Package for Comparing Biological Themes Among Gene Clusters. OMICS: A Journal of Integrative Biology 16, 284– 287 (2012).

91. 91. Csárdi, G., et al. igraph for R: R interface of the igraph library for graph theory and network analysis. Zenodo 10.5281/zenodo.10681749 (2024).

92. Shannon, P. et al. Cytoscape: A Software Environment for Integrated Models of Biomolecular Interaction Networks. Genome Res 13, 2498–2504 (2003).

93. Rohart, F., Gautier, B., Singh, A. & Cao, K.-A. L. mixOmics: An R package for ‘omics feature selection and multiple data integration. PLOS Computational Biology 13, e1005752 (2017).

94. Vertesi, A. et al. Standardized Mini-Mental State Examination. Use and interpretation. Can Fam Physician 47, 2018–2023 (2001).

